# CREaTor: Zero-shot *cis*-regulatory pattern modeling with attention mechanisms

**DOI:** 10.1101/2023.03.28.534267

**Authors:** Yongge Li, Fusong Ju, Zhiyuan Chen, Yiming Qu, Huanhuan Xia, Liang He, Lijun Wu, Jianwei Zhu, Bin Shao, Pan Deng

**Author notes:** These authors contributed equally: Yongge Li, Fusong Ju, and Zhiyuan Chen.

## Abstract

Linking *cis*-regulatory sequences to target genes has been a long-standing challenge. In this study, we introduce CREaTor, an attention-based deep neural network designed to model *cis*-regulatory patterns for genomic elements up to 2Mb from target genes. Coupled with a training strategy that predicts gene expression from flanking candidate *cis*-regulatory elements (cCREs), CREaTor can model cell type-specific *cis*-regulatory patterns in new cell types without prior knowledge of cCRE-gene interactions or additional training. The zero-shot modeling capability, combined with the use of RNA-seq and ChIP-seq data only, allows for the readily generalization of CREaTor to a broad range of cell types. Evaluation reveals that CREaTor outperforms existing methods in capturing cCRE-gene interactions across various distance ranges in held-out cell types. Further analysis indicates that the superior performance of CREaTor can be attributed to its capacity to model regulatory interactions at multiple levels, including the higher-order genome organizations that govern cCRE activities as well as cCRE-gene interactions. Collectively, our findings highlight CREaTor as a powerful tool for systematically investigating *cis*-regulatory programs across various cell types, both in normal developmental processes and disease-associated contexts.

## Background

Cell type-specific *cis-*regulatory programs allow specialized gene expression and cellular functions in eukaryotic organisms during development and differentiation (1–3). Mutations in *cis*-regulatory elements (CREs), though having no impact on protein sequences, contribute to various diseases by disrupting the normal functionality of their target genes (4–9). Decoding how CREs regulate gene expression coordinately in different cell types may reveal the mechanisms of cell identity maintenance and hint at the origins of developmental defects and human diseases.

However, linking candidate CREs (cCREs) to genes remains a substantial challenge. Experimental assays such as Hi-C (10), capture Hi-C (11), and ChIA-PET (12) have been deployed for cCRE-gene mapping, yet they measure physical proximities between elements and genes instead of direct regulatory activities. Systematic evaluation of activities of enhancers, a major type of CREs, becomes possible with CRISPR perturbation tools most recently (13–15), but only a subset of enhancers can be evaluated due to the great number of candidate enhancers in the genome (16,17). Meanwhile, the evaluations are restricted to the cell types examined in the studies.

Computational-based data-driven methods have been proposed for enhancer regulation prediction (18–22), but their performance and generalization ability are subject to limited data on *bona fide* enhancer-gene interactions, the varying number of candidate enhancers for different genes, and the complex nature of enhancer-gene regulation (7,23–25). Lack of native context during modeling is another drawback commonly seen due to a trade-off for computational feasibility. A typical case is the binary classification task where a sequence pair of enhancer and promoter is given as input (18–20), which may only recover regulation relationships mediated by conserved transcription factors universally present.

Interestingly, modeling studies on gene expression prediction from local genome sequences showed that cCRE-gene interactions were implied in the designed neural network architecture (26–28), suggesting an alternative approach for enhancer activity modeling. However, since different cell types in the same organism share the same reference genome, the sequence-based models such as Basenji2 (26) and Enformer (27), which take reference genome as input, cannot predict the activities of cell type-specific *cis*-regulatory sequences in cell types unseen by the model. While GraphReg (28) introduced a model architecture that can be generalized to new cell types, the model’s dependency on 3D genomic data narrows its applicability, as such data is not readily available.

To model cCRE-gene interactions and discover universal *cis*-regulatory patterns across cell types, we developed a hierarchical deep learning model based on the self-attention mechanism. The model named CREaTor (*Cis*-Regulatory Element auto Translator) utilizes cCREs in open chromatin regions identified by Encyclopedia of DNA Elements (ENCODE) together with ChIP-seqs of transcription factors and histone modifications (16,29) to predict the expression level of target genes. In the design, attention blocks serve as key components for accurate expression prediction, which is achieved by learning relationships between input cCREs and genes, as well as cCREs and cCREs, during training. Therefore, leveraging attention mechanisms and training on richly labeled data generated through standardized experiments, we are able to model element interactions with a zero-shot setting. In other words, the model can present cCRE-gene interactions without requiring training on such data. Moreover, since CREaTor uses cCRE landscape and ChIP-seq profiles as input, which differ between cell types, it can model CRE-gene interactions in new cell types without additional training. Using dispersed elements instead of the entire genomic context flanking each gene also greatly reduces computational costs for modeling. Testing on a held-out cell line, we show that CREaTor can effectively model the interactions between *cis*-regulatory sequences and target genes for accurate gene expression prediction. Further analysis indicates that CREaTor learns higher-order genome organization and cross-cell type regulatory mechanisms, which might explain its exceptional performance in cell types unseen by the model.

## Results

### CREaTor predicts cell type-specific gene expression in unseen cell types

CREaTor consists of two transformer models at different resolutions (Fig. 1a and Extended Data Fig. 1). Transformer is a deep learning architecture that has been demonstrated as a powerful tool for natural language processing (30–32), computer vision (33,34), and biological modeling (27,35,36). A core component of a transformer is the self-attention module, which extracts sequence-level information by modeling the interactions between elements at different positions in the sequence (30). In CREaTor, the lower-level transformer (element encoder) learns the latent representation for each cCRE from the DNA sequence and chromatin states of the element itself, while the upper-level transformer (regulation encoder) predicts gene expression from a collection of cCRE latent representations flanking the target gene. Self-attention extracted from the regulation encoder is used to interpret the cCRE-gene and cCRE-cCRE interactions.

**Figure 1.**
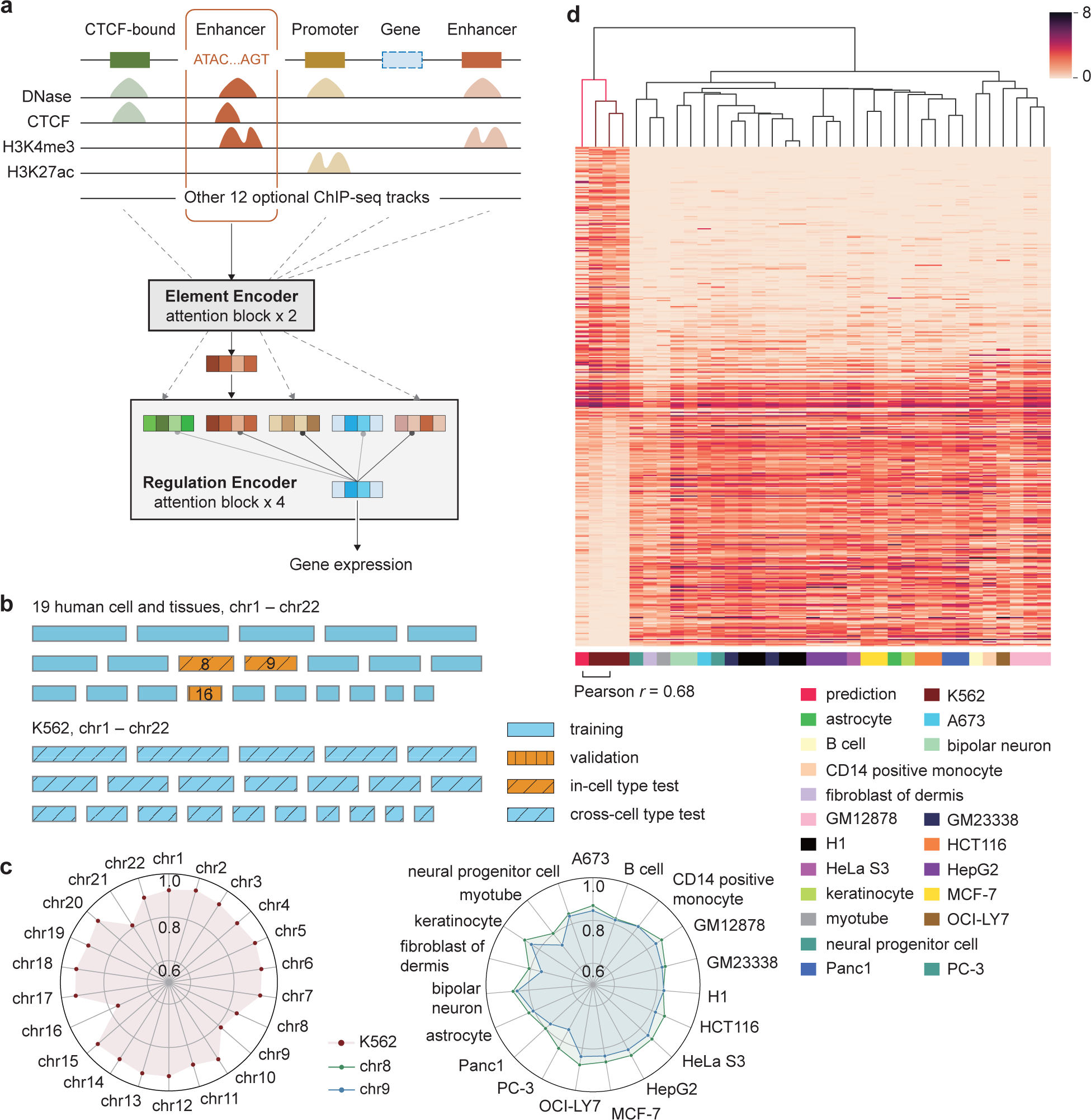
Accurate gene expression prediction with CREaTor. **a)** Schema of CREaTor. The model predicts target gene expression from the flanking cCREs with a hierarchical transformer structure. Localization of cCREs was obtained from ENCODE consortium. A combination of genomic sequences, chromatin openness, and a collection (3–13) of ChIP-seq profiles was used as input features for each cCRE. **b)** Visualization of data split strategy: we trained our model on gene expression of 19 autosomes from 19 different cell lines and tissues respectively. Genes on chr16 from the 19 cell lines and tissues were used for parameter tuning (validation), while genes on chr8, 9 were used for model evaluation (in-cell type test chromosomes). Genes from all autosomes in K562 (cross-cell type test chromosomes) were detailedly evaluated to demonstrate the model’s ability on cross-cell type gene expression and regulation modeling. Also see in Supplementary Figure 1. **c)** Pearson *r* between observed and predicted expression of genes. Left: Pearson *r* between observed and predicted expressions of genes on cross-cell type test chromosomes. Right: Pearson *r* between observed and predicted expressions of genes on in-cell type test chromosomes. Green and blue dots indicate chr8 and 9 respectively. See Extended Data Table 2 for results with different random seeds. **d)** Clustering map of predicted and observed expression of K562 specific genes (calculated with RSME, see Methods) in different cell types. The leftmost column is the predicted value, which is clustered with the K562 observed gene expression data using the hierarchical clustering method. Expression values were transformed with log1p. Observed gene expression profiles from different sources (with different experiment IDs on ENCODE) for the same cell type are calculated independently.

We trained CREaTor on 19 human tissues and cell lines whose annotated cCRE information was available in SCREEN Registry (16) (Supplementary Table 1). In each cell type, chr16 was left out for validation, chr8 and 9 were left out for testing (*in-cell type test chromosomes*), and all other autosomes were used for training (Fig. 1b and Supplementary Figure 1). By training the model with data from multiple cell types jointly, we expected the model to learn general rules guiding gene regulation across cell types. Next, we evaluated CREaTor’s performance on autosomes of the K562 cell line (*cross-cell type test chromosomes*), which were unseen by the model, to demonstrate the generalizability of our method (Fig.1b and Supplementary Figure 1). For the in-cell type test chromosomes, CREaTor reached a mean correlation of 0.850 and 0.818 (Pearson *r*) for chr8 and 9 respectively (Fig.1c and Extended Data Table 1). While for the cross-cell type test chromosomes, the correlations between observed and predicted gene expression on different chromosomes ranged from 0.756 to 0.936, with a mean correlation of 0.902 (Pearson *r*) (Fig. 1c and Extended Data Table 2). Notably, the predictive accuracy of K562 chr8 and 9 (0.839 and 0.810 respectively, Pearson *r*) was comparable to that of in-cell type test chromosomes, suggesting that CREaTor can predict gene expression efficiently from cCREs in new cell types.

However, the performance gap between chr8/9 and other chromosomes in K562 is non-trivial. We reasoned that the presence of housekeeping genes and several hematopoietic cell types in training data alleviated the challenge for expression prediction on chromosomes other than 8 and 9. To assess the generalizability of CREaTor more rigorously, we next examined if CREaTor could make cell type-specific predictions. With gene differential expression (GDE) analysis on paired data between K562 and each of the 19 cell types used for model training respectively, we identified 410 genes that were differentially expressed in the K562 cell line (Methods). For a number of these genes, including hematopoietic regulators KLF1 and TAL1 and hemoglobin subunit protein HBE1, CREaTor made a prediction on rival with experimental quantifications (Extended Data Fig. 2). In addition, single-linkage clustering analysis demonstrated that the prediction on K562 differentially expressed genes was more similar to observed K562 expression compared to other cell types (Pearson *r*=0.68, Fig. 1d).

To further demonstrate that the accurate prediction of K562 expression is not attributed to the similarity between K562 and training cell types, we compared the predicted expressions with 122 observed expression profiles of 20 distinct cell types. These profiles included 12 profiles from 3 independent K562 RNA-seq experiments that our model had not previously encountered. We visualized all expression profiles with Uniform Manifold Approximation and Projection (UMAP) in a 2-dimensional space. For different chromosome subsets, predicted expression consistently exhibits high similarity to K562, as opposed to other cell types, including those sharing hematopoietic origins with K562 (Supplementary Figure 2). Also, we conducted leave-one-chromosome-out and leave-one-cell type-out experiments to confirm that CREaTor’s superior performance was not limited to chr8-9 and K562, respectively (Supplementary Figure 3 and Supplementary Table 2).

It has been reported that histone modifications and DNA openness proximal to gene transcription start sites (TSS) are significantly correlated with active transcription (37). To demonstrate that distal information contributes to model performance, we compared models trained with cCREs up to 2kb, 5kb, 10kb, 100kb, or 1Mb away from the TSS of target genes. Performance improved with increasing candidate window sizes (Extended Data Fig. 3), suggesting that CREaTor predicts gene expression from both proximal and distal cCREs. Also, this result indicates that distal cCREs are substantial for accurate expression prediction, supporting the importance of long-range *cis*-regulatory interactions in gene regulation. But meanwhile, it is worth noting that the model trained with cCREs up to 2kb away from target genes performed significantly better than random guesses, consistent with the knowledge that the proximal functional genomes and cCREs are closely related to gene expression.

### Self-attention reveals functional cCREs in unseen cell types

Attention weights between cCREs and target genes extracted from CREaTor (Methods) may be exploited to interpret the importance of each cCRE to genes. To test this hypothesis, we benchmarked CREaTor against 3 CRISPR-based experimental-validated K562 enhancer-target gene datasets (13–15). To be noted, criteria for candidate enhancers vary in each study and few enhancer-target gene pairs tested were shared among studies (Extended Data Fig. 4 and Supplementary Table 3). Thus, we combined the experimental results and identified 1859 putative enhancers related to 328 genes that were tested by both the experimental approaches and CREaTor across the K562 genome. CREaTor prioritizes positive enhancer-gene pairs to negative ones with larger attention scores (auROC=0.834, auPRC=0.620; Fig. 2a-b) and the performance is further improved when we adjusted the attention scores with enhancer-gene genomic distances (auROC=0.843, auPRC=0.667; Fig. 2a-b). In addition, we compared the scores derived from the attention weights of CREaTor with a quantitative analysis of enhancer effects as described in a previous study (13). In this study, the enhancer effect on gene expression was defined as the change in gene expression upon enhancer knockdown using CRISPR perturbation. Consequently, the quantitative effect is inversely related to the enhancer activity. In line with this understanding, we observed a negative correlation, with a Spearman *ρ* of -0.269, between the CREaTor scores and the quantitative observations (Fig. 2c), implying that CREaTor captures quantitative effects of cCREs to genes.

**Figure 2.**
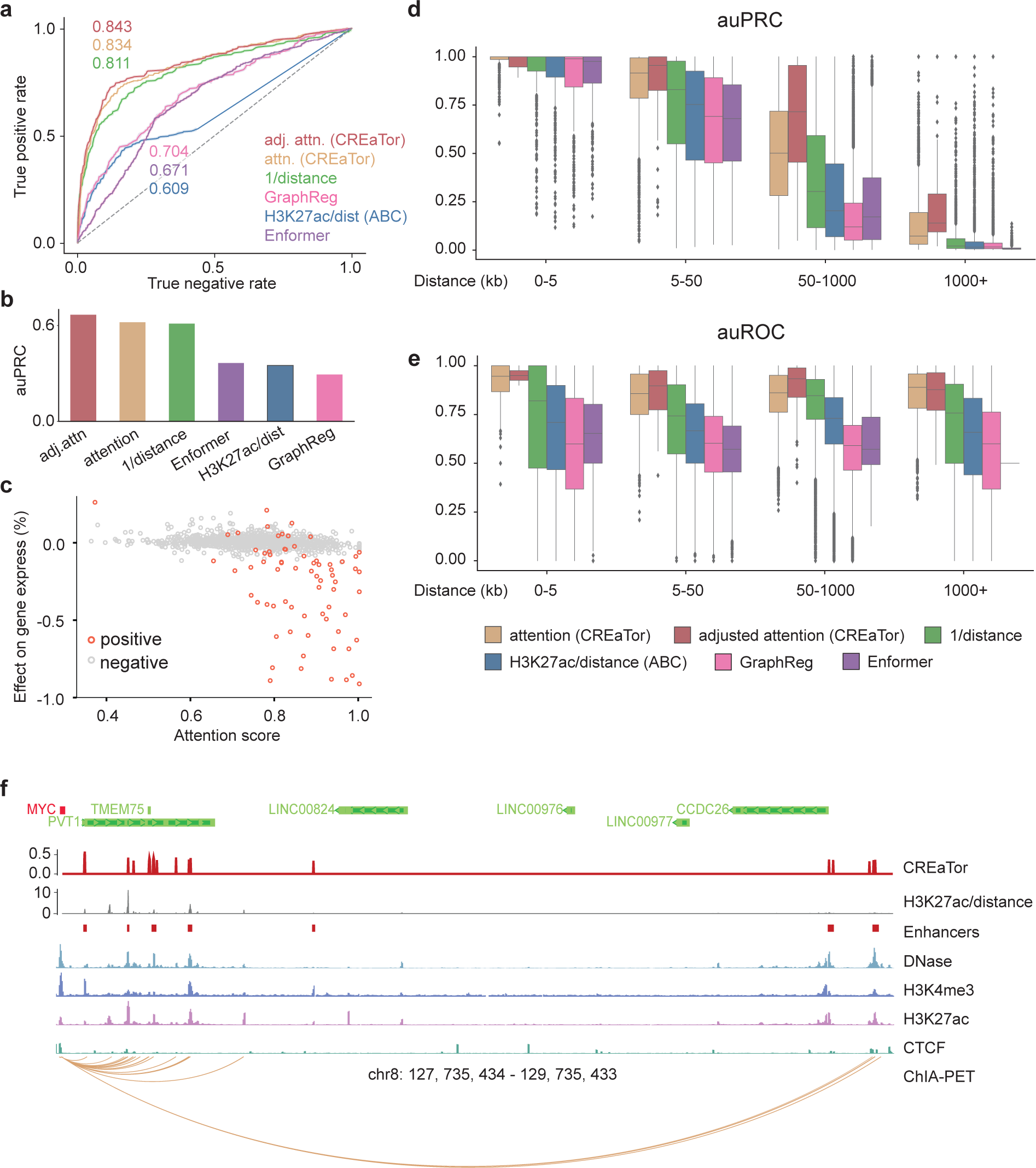
Attention matrix of CREaTor implies cCRE-gene interactions. **a-b)** auROC (a) and auPRC (b) of CREaTor outperform its counterparts on cCRE-gene pair classification. Attention (attn., yellow): normalized attention weights (genes to cCREs) in CREaTor. Adjusted attention (adj. attn., red): attention scores / log10 (distance). H3K27ac/dist (blue): approximate of the ABC score. Distance quantifies relative genomic distances between genes and cCREs. H3K27ac value of a cCRE is calculated as the sum of H3K27ac peak values of the element. Labels (positive/negative) of cCRE-gene pairs were collected from 3 independent CRISPR perturbation experiments. **c)** Attention scores derived from attention weights are significantly correlated with the effect of enhancer on gene expression quantified by Fulco et al^13^. As the quantification measures the change of target gene expression upon enhancer knock-down using CRISPR perturbation, the quantitative effect values are invertsely related to enhancer activities. **d-e)** auPRC (d) and auROC (e) of CREaTor and its counterparts on the classification of cCRE-gene pairs collected from a Pol-II mediated ChIA-PET experiment. The performance is evaluated for each gene and each distance group separately. Groups with <10 samples were filtered out. Center line, median; box limits, upper and lower quartiles; whiskers, 1.5x interquartile range; points, outliers. **f)** MYC locus showing predicted and previously reported regulators in K562 cells. For CREaTor (red) and H3K27ac/distance (gray), peaks on the tracks represent the scores of different cCRE regions. Enhancers track (red squares) denotes reported active regulators of MYC. Representative DNase, H3K4me3, H3K27ac and CTCF tracks as well as ChIA-PET interactions in K562, are also annotated.

We also compared CREaTor with 4 methods previously used for cCRE-gene interaction modeling: 1) Predictions based solely on genomic distances between cCREs and genes; 2) Predictions based on cCRE H3K27ac signals and cCRE-gene distances (approximate version of the Activity-by-Contact (ABC) score(13)). These model-free approaches can estimate activities of cCREs spanning varying ranges without prior knowledge of *cis*-regulatory programs in any cell types, or cell types with H3K27ac quantifications, which align well with the setting of CREaTor. Evaluated on a comprehensive set of metrics, CREaTor outperforms both methods at different distance groups (Fig. 2a-b and Extended Data Fig. 5). In addition, we compared CREaTor to 2 state-of-the-art deep learning approaches, 3) Enformer (27) and 4) GraphReg (28). Both Enformer and GraphReg, trained with supervised gene expression prediction tasks, support zero-shot cCRE-gene interaction prediction. However, Enformer’s architecture limits it from long-range enhancergene interaction prediction, as the released Enformer model can only predict interactions up to 200kb. Additionally, it cannot generalize to new cell types as it solely relies on genomic sequences for predictions. To simulate prediction tasks in new cell types, we adopted the cell-type-agnostic setting of Enformer (Methods). As expected, predicting enhancer-gene interactions in new cell types with Enformer is not favorable (Fig. 2a-b and Extended Data Fig. 5). GraphReg, on the other hand, predicts CAGE signals from 1D epigenomic data and 3D genomic structures, allowing it to generalize to new cell types. However, its dependency on 3D genomic structures and CAGE profiles narrows its applicability. To evaluate GraphReg, we trained an enhanced GraphReg model using 9 cell types and 16 types of epigenomic profiles from scratch and derived feature importance to estimate enhancer activities in K562 as suggested by the original study (Methods). Our results show that CREaTor greatly outperforms GraphReg (Fig. 2a-b and Extended Data Fig. 5), suggesting the superiority of CREaTor’s design.

Next, cCRE-gene interactions discovered by CREaTor were further benchmarked against a genome-wide Pol II-mediated ChIA-PET dataset (38). Compared with CRISPR perturbation studies, ChIA-PET covers a broader range of genes and regulators, thus capturing more comprehensive interactions between genes and regulators. We recovered 6, 132, 740 cCRE-gene pairs (both positive and negative) across the K562 genome from ChIA-PET. To benchmark CREaTor and its counterparts, for each gene, we calculated auROC and auPRC of the corresponding cCRE-gene pairs stratified by their relative genomic distances. Among all, CREaTor shows the highest median auROC and auPRC for gene collections at all distance groups and greatly outperforms Enformer and GraphReg (Fig 2d-e). Strikingly, CREaTor performs substantially better at groups spanning longer ranges.

Since ChIA-PET captures physical proximities between genomic regions, false positives exist when active CRE-gene pairs are recovered from ChIA-PET. To benchmark our method more comprehensively, we calculated the precision and specificity scores for different methods considering that these metrics are less impacted by false positives. Consistently, CREaTor outperforms other methods (Extended Data Fig. 6), indicating that CREaTor can capture cCRE-gene interactions efficiently from genomic features flanking target genes in unseen cell types.

Lastly, we examined if our model recovered regulators of the oncogenic gene MYC (chr8: 127,735,434-127,742,951). cCREs of MYC disperse along genomic sequences to as far as 2 Mb downstream MYC TSS and active MYC regulators in K562 were identified by previous studies with various approaches(39–41). Therefore, we examined if CREaTor could pinpoint these regulators accurately. The result indicates that CREaTor prioritizes positive MYC cCREs with larger attention scores and captures active cCREs missed by other predictive approaches (Fig. 2f). In addition, 2 groups of sharp peaks are observed 2Mb downstream MYC TSS (Fig. 2f), in concordance with the existence of 2 distal super-enhancer regions of MYC. Since MYC is in both in- and cross-cell type test sets, we believe that CREaTor has learned general rules guiding cCRE-gene interactions in different cell types, rendering it an efficient tool for cCRE activity modeling in unseen cell types.

### CREaTor captures chromatin domain boundaries in unseen cell types

Three-dimensional (3D) chromatin folding allows physical interactions between distal cCRE and genes and the information can also guide gene regulation modeling (13,28,42). Without incorporating 3D chromatin folding information in our model, we were curious to see if CREaTor captured the topological structure of the genome, considering that CREaTor precisely recovers cCRE-gene interactions even of long ranges.

Attention matrices extracted from the model imply not only the interactions between cCREs and genes, but also relationships between cCRE-cCRE pairs. To examine if the attention matrix reflects contact frequency between elements, we aggregated the attention matrix at 10kb resolution for each gene in K562 and compared the results to a high-resolution Hi-C study (10). In addition to observing similar checkerboard patterns between the attention matrix and Hi-C (Fig. 3a), we systematically evaluated the consistency between the attention matrix and topologically associating domains (TADs) by analyzing insulation scores (Methods). We calculated insulation scores from the attention matrix over 12,584 K562 TAD boundaries defined in a recent study (43). The average score across the genome shows a clear insulation pattern on boundaries, similar to that calculated from the Hi-C experiment (Fig. 3b). Meanwhile, no significant decrease over GM12878-specific TAD boundaries (43) is observed with insulation scores calculated from either K562 attention matrix or K562 Hi-C (Fig. 3b), demonstrating that CREaTor captures K562-specific TAD boundaries. Together, the results show that CREaTor can infer cell type-specific topological structures of genomes in cells unseen by the model.

**Figure 3.**
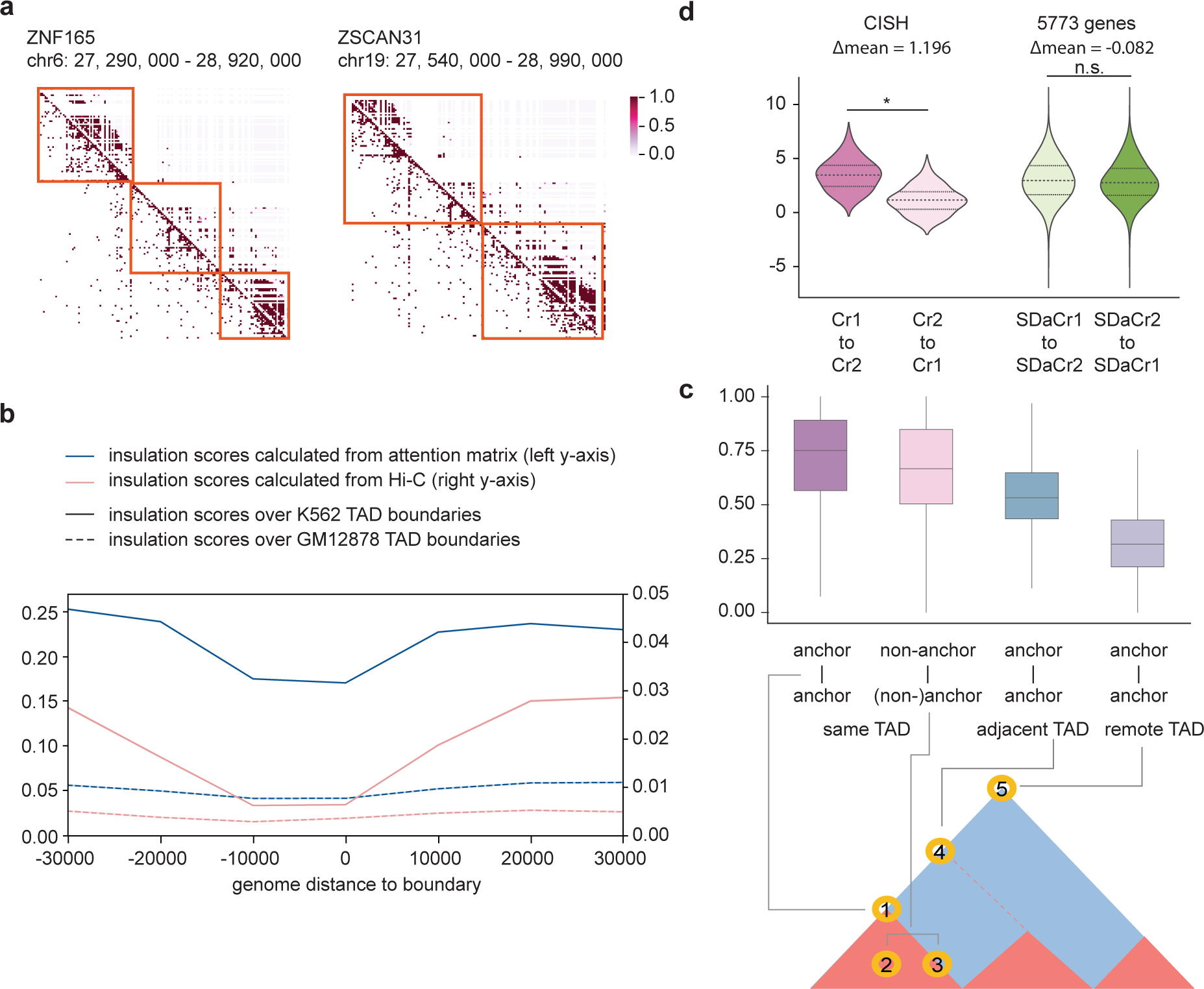
CREaTor captures hierarchically higher-order genome organizations. **a)** Example genomic regions showing the similarity between attention matrix (above the diagnal) and Hi-C contact matrix (below the diagonal). Orange boxes indicate TAD domains. **b)** Average insulation scores across the K562 genome at 10-kb resolution calculated from attention matrix and Hi-C. Blue line and left y-axis: insulation scores of attention matrix. Pink line and right y-axis: the insulation scores of Hi-C. Solid lines indicate insulation scores over K562 TAD boundaries and dashed lines indicate insulation scores over GM12878 boundaries. The x-axis is centered on TAD boundaries. **c)** Upper panel: Statistics of attention weights between CTCF-bound element pairs with different topological relationships. Center line, median; box limits, upper and lower quartiles; whiskers, 1.5x interquartile range. Lower panel: illustration of CTCF-bound element pairs used for the analysis. The red triangle represents TAD domains called from the Hi-C matrix (blue). **d)** Average attention scores between elements without normalization. p-value is calculated with Mann-Whitney U Test.

We reason that CREaTor infers genome structures by learning the insulating behaviors of CTCF-bound elements. Consistently, we found that paired CTCF-bound insulators flanking the same TAD domain showed significantly larger attention scores compared to either unmatched insulator pairs spanning multiple TADs, or pairs involving non-insulator CTCF-bound elements (Fig. 3c). Thus, CREaTor may predict gene expression and capture cCRE-gene regulation efficiently by modeling topological patterns of the genome.

### CREaTor implies directional regulation between cCREs

It is long proposed that enhancers form hierarchical relationships with each other, yet the relationship is challenging to be disentangled with biological experiments. For example, Carleton et al. developed an enhancer interference technique (Enhancer-i) to study the combinational effects of distal regulatory regions on genes (44). They showed the interdependence between CISH-1 and CISH-2, two estrogen receptor α-bound enhancers of the cytokine signaling suppressor gene *CISH*. However, the detailed mechanism between interactions of CISH-1 and CISH-2 could not be elucidated. Here, we examined attention scores between cCREs within CISH-1 and CISH-2 regions (denoted as Cr1 and Cr2 respectively) and found that the attention from Cr1 to Cr2 is significantly larger than the other way around (Fig.3f). To rule out potential distance bias, we examined attention score distribution of 5773 genes whose cCRE-gene distances were similar to Cr1-*CISH* and Cr2-*CISH* (denoted as SDaCr1 and SdaCr2). Remarkably, no directional preference between SDaCr2 and SDaCr2 was observed (Fig.3f). Therefore, our results indicate that there could be a directional relationship between CISH1 and CISH2, which is driven by hierarchical regulation of enhancers. We thus believe that with further development, CREaTor has the potential to become a powerful tool for understanding the causal relationships within enhancer networks.

### cCRE representations learned by CREaTor suggest a new role of CTCF-bound elements

To investigate how CREaTor perceives cCREs and their roles during gene regulation, we clustered cCREs by the 256-dimensional cCRE representations extracted from CREaTor and examined features enriched in each group.

While different cCRE types are enriched in different clusters (Fig. 4a-b), cCRE representations learned by our model better capture functional variations of elements compared to the classification of ENCODE. For instance, while cCREs are aggregated into 6 clusters, both cluster 0 and cluster 1 are enriched with proximal enhancer-like elements, cCREs that show enhancer-like signatures falling within 200bp of an annotated transcript start site (TSS) (16). However, proximal enhancer-like elements in cluster 0 are enriched for RNA polymerase II (Pol II) signals (Extended Fig. 7), markers of active transcription events, compared to those in cluster 1. Since promoter-like elements are also enriched in cluster 0 and enhancers are believed to be able to contribute to promoter activities (45), we reason that CREaTor learns the discrepancies between enhancer-like elements of different roles and therefore associates a subgroup of proximal enhancer-like elements with promoters. Meanwhile, the fuzzy boundaries between clusters may indicate the adaptable functions of elements for gene regulation captured by our model.

**Figure 4.**
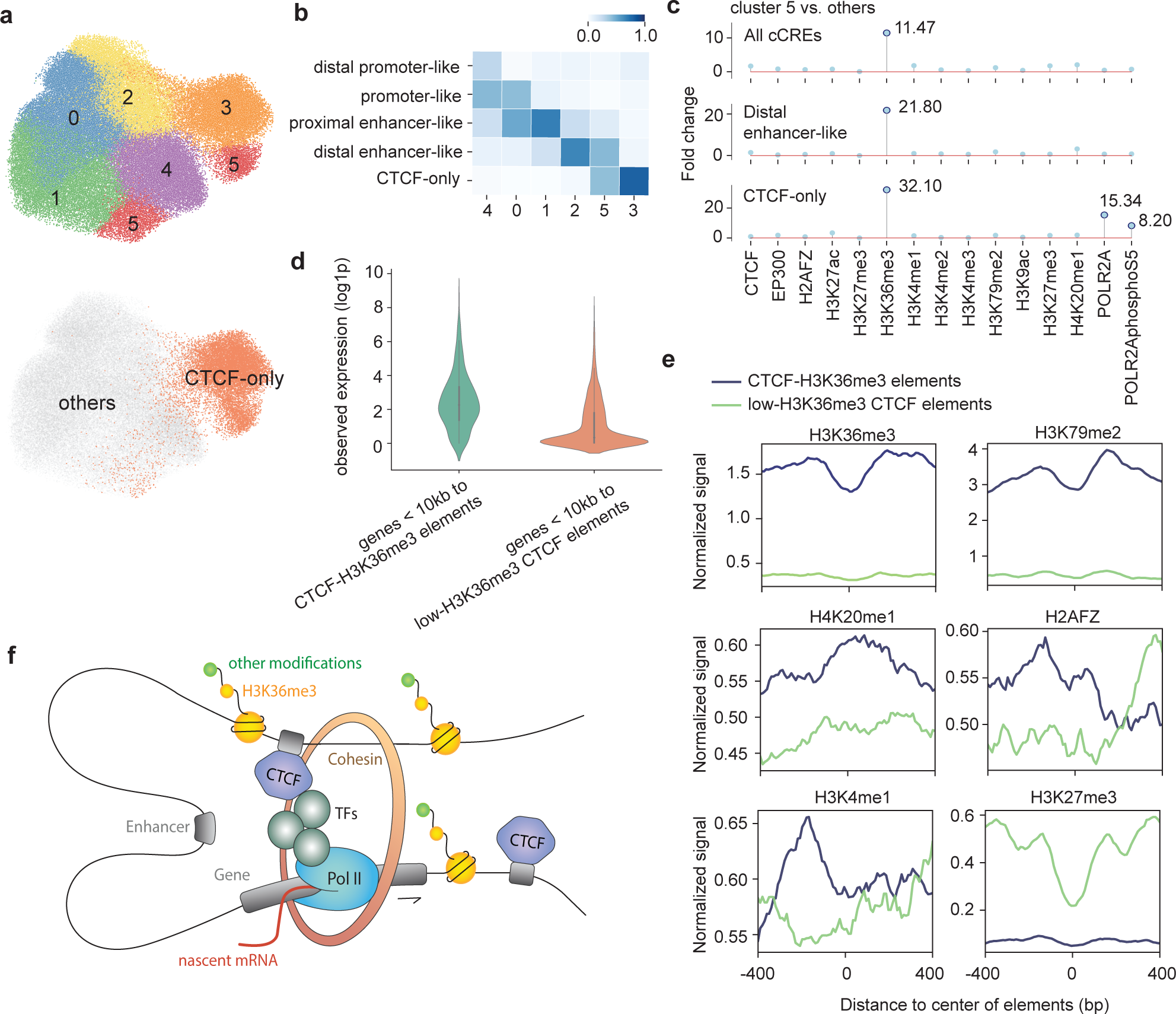
cCRE representations learned by CREaTor suggest a new role of CTCF-bound elements. **a)** Uniform Manifold Approximation and Projection (UMAP) of cCRE embeddings in K562. Upper: colored and numbered as clusters grouped by the Leiden algorithm. Bottom: colored and labeled by element type. **b)** Composition of different element types in each cluster by percentage. Proximal elements: elements falling within 200bp of an annotated TSS. Distal elements: elements more than 200bp from any annotated TSS. Promoter-like: elements with high DNase and H3K4me3 signals. Enhancer-like: elements with high DNase and H3K27ac signals. CTCF-only: elements with high DNase and CTCF signals, as well as low H3K4me3 and H3K27ac signals. **c)** Fold change of histone marker peaks of given types of cCREs in cluster 5 with respect to those in other clusters. Top: all cCREs. Middle: distal enhancer-like elements. Bottom: CTCF-only elements. **d)** Expression value (log1p) distribution of genes within 10kb of different types of CTCF-bound elements. **e)** Average signals of H3K36me3, H3K79me2, H4K20me1, H2AFZ, H3K4me1 and H3K27me3 on different types of CTCF-bound elements. **f)** Illustration for the proposed model of CTCF-H3K36me3 elements promoting transcription elongation.

CTCF-only cCREs, which lack both enhancer-like signatures and promoter-like signatures, are more isolated from other elements, consistent with their insulator and looping functions (Fig. 4a). However, CTCF-only cCREs are clustered into 2 separate groups, while a subgroup of CTCF-only cCREs is aggregated with distal enhancer-like elements in cluster 5 (Fig. 4b). Compared to other clusters, cluster 5 shows a significant enrichment of H3K36me3 peaks (Fig. 4c), a histone modification associated with diverse functions in conjugation with different types of epigenetic markers (42,46–50), indicating a higher chromatin activity of these elements. Consistent with the result, genes close to CTCF-only cCREs in cluster 5 (denoted as CTCF-H3K36me3 elements) show higher expression values compared to those close to low H3K36me3 CTCF-only elements (Fig. 4d), suggesting a more active role in gene transcription of CTCF-H3K36me3 elements.

Depletion of repressive histone modification H3K27me3 also supports the greater activity of CTCF-H3K36me3 elements (Fig. 4e). Other from H3K36me3, CTCF-H3K36me3 elements are enriched with H3K79me2 and H4K20me1 (Fig. 4e), a pattern that has been previously reported to be associated with active transcription and splicing of exons(46). Meanwhile, CTCF-H3K36me3 elements show increased H3K4me1 and H2AFZ signals (Fig. 4e), both of which are associated with enhanced transcription elongation(51,52). Considering a majority of CTCF-H3K36me3 elements locate outside exon regions, we propose that CTCF-H3K36me3 elements promote transcription elongation by serving as binding hubs for various *cis*- and *trans*-regulatory elements (Fig. 4f), which are captured by CREaTor for cross-cell type gene regulation modeling.

## Discussion

While profiling gene expressions and epigenetic modifications in various cell types is feasible, systematical approaches profiling cell type-specific *cis*-regulatory patterns are currently not achievable. As a result, deep learning techniques, despite greatly advancing our understanding of gene regulation in many areas, face challenges in this area due to the lack of training data. To overcome this challenge, we introduce the CREaTor framework. By strategically selecting training tasks and incorporating attention mechanism, CREaTor enables zero-shot *cis*-regulatory pattern modeling and cCRE-gene interaction prediction at ultra-long range. In addition, it can generalize to new cell types without requiring additional training or relying on 3D genomic data, making CREaTor versatile and applicable to a wide variety of cell types.

Comprehensive validation and benchmark experiments show that our model outperforms alternative methods in modeling cCRE-gene interactions. Additionally, attention analysis shows that CREaTor learns cell type-specific 3D genome interactions and insulation behaviors, which play crucial roles in gene regulation, during gene expression prediction. These results indicate that our model is able to capture the underlying principles that guide cCRE-gene interactions across different types of cells, utilizing 1D features such as histone modifications on the genome. Further experiments showcase that CREaTor captures regulatory mechanisms at multiple levels. Aside from cCREs, CREaTor also learns gene interpretations during modeling. Our model stratifies genes into distinct groups enriched with different biological processes and molecular functions (Extended Data Fig. 8), indicating that CREaTor has captured active pathways mediated by different transcription factor programs, which allow cell type-specific gene regulation by binding to cCREs. These analyses may explain how our model captures *cis*-regulatory patterns from a range of cross-cell type gene expression predictions.

Except for modeling cross-cell type *cis*-regulatory patterns, the adoption of transformer architecture has allowed for greater flexibility during application. For instance, the element module in CREaTor can handle candidate regulators of different lengths. Also, the regulation module allows the modeling of gene context with varying numbers of cCREs spanning varying genomic ranges. In addition, despite 17 types of input features being used for training, our model can still predict gene expression and infer cCRE-gene interactions when some features are missing, though a lack of features may negatively impact the performance of the model (Fig. 5a). Overall, this flexibility makes CREaTor more adaptable to different situations compared to other methods.

**Figure 5.**
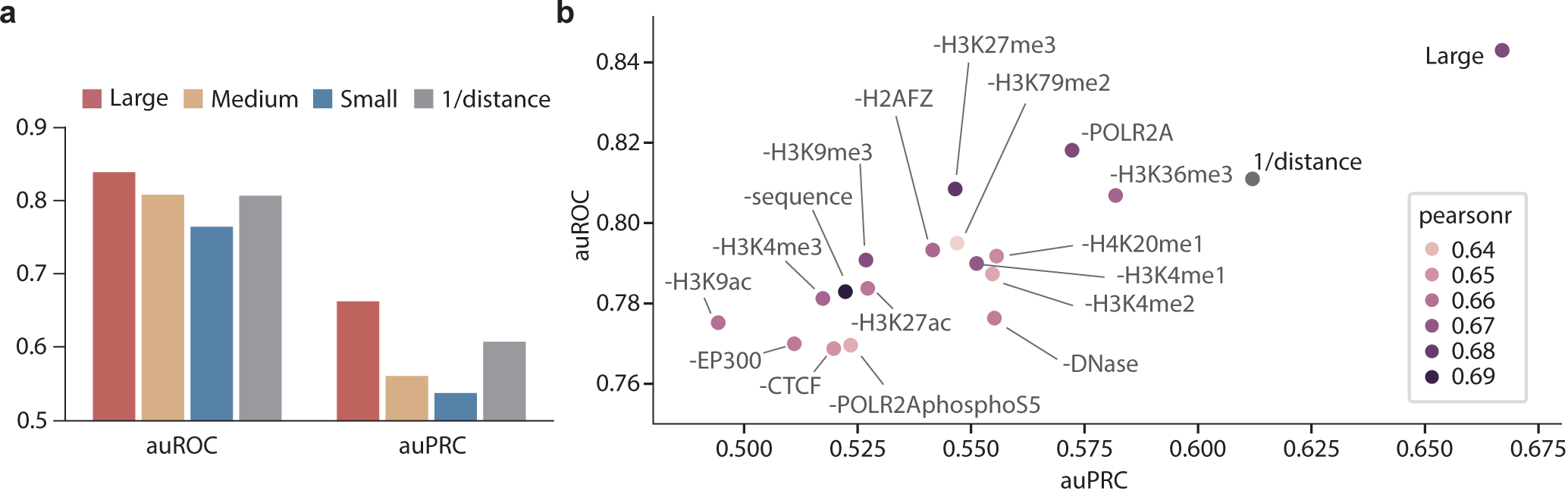
Feature ablation study demonstrates the importance of feature integration for modeling. **a)** auROC and auPRC of 4 different models on cCRE-gene pair classification. Large (red): the model trained with 17 types of features. Medium (yellow): the model trained with 8 types of features (genomic sequence, DNase, CTCF, H3K27ac, H3K4me3, H3K9ac, EP300, and POLR2AphosphoS5). Small (blue): the model trained with 5 types of features (genomic sequence, DNase, CTCF, H3K27ac, and H3K4me3). **b)** Large model trained with 17 types of features outperforms other models on cCRE-gene interaction classification tasks. Minus signs indicate the following type of feature is removed during model training. Labels (positive/negative) of cCRE-gene pairs were from the same source as Figure 2. The colors of dots indicate the Pearson r between observed and predicted expression of K562-specific genes.

In order to assess the impact of each input feature on predicting gene expression and modeling *cis*-regulatory patterns, we conducted an ablation study by excluding individual feature types from the model’s training. Our results revealed inconsistent performance between different tasks - while genome sequence information is dispensable for successful cell type-specific gene expression prediction, it has a moderate impact on the accuracy of CRE-gene interaction inference (Fig. 5b). Likely due to complementary relationships between different feature types, no single feature was found playing a dominant role in CRE-gene interactions. However, the exclusion of any feature type leads to decreased performance for CRE-gene interaction inference, and the model trained with a full collection of features performs significantly better on the cCRE-gene interaction classification task compared to all other settings (Fig. 5b), suggesting that utilization of multiple types of features guarantees our model’s performance across cell types and CREaTor may have learned synergistic relationships between features for accurate *cis*-regulatory pattern modeling. Among all, features that are known to be crucial for gene regulation, such as CTCF, DNase, H3K4me3, H3K27ac, H3K9ac and EP300 show greater importance. Pol II with enriched phosphorylated Ser5 in CTD is more important for gene expression prediction and cCRE-gene interaction inference than its unphosphorylated form. This could be explained by the active involvement of phosphorylated CTD in binding *trans*- and *cis*-regulatory elements for dynamic transcription regulation. Evaluating the impact of different Pol II phosphorylation states on gene regulation modeling in the future might give additional insight into their roles. Interestingly, the results imply a paradoxical role of H3K36me3 in gene regulation. This may be due to the fact that the gene sets regulated by H3K36me3 are not included in the CRISPR perturbation experiments.

It is worth pointing out that our model’s performance is constrained by the limited accessibility of functional genomic data, regardless of the features employed. Although the ENCODE project provides various high-quality functional genomic data of many cell types, the coverage is still limited due to the vast number of cell types, histone modifications, and proteins binding to the genome. For example, cohesin, which regulates chromatin structure by participating in the loop extrusion process, was not included in our model data at the time of modeling due to the lack of data in most cell types. We believe that incorporating such data would further improve the generalizability of our method.

Compared to previous approaches, CREaTor is able to capture distal *cis*-regulatory patterns and infer cCRE-gene interactions spanning ultra-long distances. We believe that one reason for this improvement is the fact that our model was trained using only cCREs. However, it is also important to note that this approach may lead to bias and neglect of atypical regulators, such as non-canonical enhancers and other low-H3K27ac regulatory elements without typical enhancer chromatin features (53,54). We expect that an end-to- end setting incorporating a deep learning module calling CREs directly from the genome will alleviate the issue of bias and allow for a more comprehensive understanding of *cis*-regulatory elements.

Finally, in the interest of simplicity and consistency with previous studies, we have chosen to utilize reference genomes during the training process. However, it is important to note that functional genomic data on ENCODE might have originated from cells with different genomes. Specifically, cell lines may exhibit different nucleotide polymorphisms, structural variations, and karyotypes. As previous studies have demonstrated the predictive capability of genomic sequences in various tasks (26,27,55,56) and we have shown that the absence of sequences negatively impacts the performance of cCRE-gene interaction inference (Fig. 5a), we anticipate that improved model performance will be garnered by considering the diverse variations and associated consequences of different cell types in future work. Despite these limitations, we believe that CREaTor can serve as a powerful tool for studying cell type-specific *cis*-regulatory patterns and gene regulation networks, with further improvements to be made in the future.

## Methods

### Model

#### Model architecture

The backbone of CREaTor is composed of two modules: (1) an element module to extract features of cCREs and (2) a regulation module to model the regulations between cCRE and genes.

CREaTor takes 200 cCREs from up- and down-stream of target gene TSS respectively as input (Note: we have also tried taking cCREs within the ± 1Mb range of a gene TSS for training. The outcomes of both strategies are comparable). Each element is represented by its DNA in the form of one-hot encoding (A = [1, 0, 0, 0, 0], T = [0, 1, 0, 0, 0], C = [0, 0, 1, 0, 0], G = [0, 0, 0, 1, 0], N = [0, 0, 0, 0, 1]) and ChIP-seq/DNase-seq with read-depth normalized signal or fold change over control, although the absence of ChIP-seqs can be tolerated by our proposed framework. We map the input DNA and ChIP-seq/DNase-seq to DNA embedding and ChIP-seq embedding through a linear projection to 256 channels respectively. Then, we organize the feature embedding at each base pair (*Emb_bp_*) as the sum of DNA embedding and ChIP-seq embedding.

The core of the element module is an element encoder based on transformer encoder architecture. Each transformer encoder layer consists of a multi-head self-attention sub-layer and a position-wise fully connected feed-forward network sub-layer^30^. In the self-attention sub-layer, scaled dot-product attentions are performed as follows: embeddings calculate the query *Q* ∈ ℝ*^nxdk^*, key **K* ∈ ℝ^nxdk^*, and value *V ∈ ℝ^nxdv^* through linear projection where *n* is the number of embeddings, *d_k_*, *d_v_* is the number of channels; the attention weight is calculated by 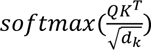 representing the attention between pairwise; lastly, the value representing the semantics of all embeddings are aggregated according to the attention weights as shown in the equations below. Feed-forward network sub-layers introduce non-linearity and interact channel information. Since the transformer encoder is a position-agnostic architecture, we apply a relative positional embedding onto the attention weights to introduce positional information. We follow T5(57) to formulate the position embedding *θ*, where *P* is the relative position between base pairs within elements.

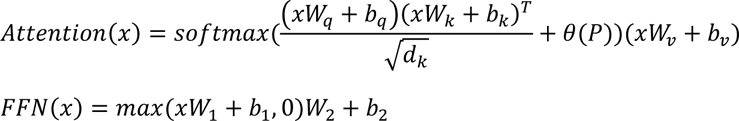

We concatenate a learnable [CLS] token to *Emb_bp_* in the element encoder. The [CLS] token adaptively attends *Emb_bp_* and we use its output as the representation of elements (*Emb_ele_*). The element encoder consists of 2 transformer encoder layers with 4 heads.

The regulation module comprises a regulation encoder to model the interactions between genes and cCREs and a prediction head for gene expression prediction.

Regulation encoder shares a similar architecture with element encoder, but with 4 transformer encoder layers and 4 attention heads. We concatenate [GENE] tokens and the corresponding cCRE embeddings *Emb_ele_* to formulate the input of regulation encoder. To be noted, [GENE] tokens are initialized by shared learnable embeddings and different genes are distinguished by their associated TSS positions. Additionally, to ensure proper information flow, we mask out the attention weight between genes. Accordingly, [GENE] tokens adaptively attend *Emb_ele_* and we use their output as the representation of genes (*Emb_gene_*). Relative position *P* is calculated as the relative genomic distances of gene TSS and elements.

At last, we apply a prediction head comprised of a linear projection and a soft plus activation to predict the gene expression given gene representations *Emb_gene_*output from the regulation encoder.

#### Model training

We trained our model with a batch size of 8 for 50, 000 steps using AdamW optimizer. For training stability, we warmed up the learning rate in the first 5, 000 steps from 0 to 1e-3 and linearly decayed it to 1e-8. Following previous work (26), we calculated the loss between the ground-truth and predicted values through a Poisson negative log-likelihood function. We also applied a gradient clip by norm with a maximum norm of 1.0 and a dropout rate of 0.1.

We verified the robustness of our model with 5 random seeds.

#### Attention score

Attention logit matrices were extracted from each attention layer in the Regulation Encoder. Both min-max and softmax normalization were applied based on needs. For cCRE-gene interaction modeling, we focused on attention from gene to cCRE only.

### Training data

RNA expression, DNase-seq, and ChIP-seq files were downloaded from ENCODE (https://www.encodeproject.org/, by October 2021). For better quality control, we used experiments included in the reference human epigenomes (29) only (ENCODE-Reference epigenome matrix). The complete list of data can be found in Supplementary Table 4.

#### RNA-seq processing

Total RNA-seq and polyA plus RNA-seq data in human biosamples were downloaded from ENCODE. Released transcript quantifications mapped to the GRCh38 sequences and annotated to GENCODE V29 were retained. Gene expression level was calculated as the sum of transcript TPM. Log1p normalization was performed.

#### DNA-seq and ChIP-seq processing

DNase-seq, histone ChIP-seq, and TF ChIP-seq files of human biosamples mapped to the GRCh38 sequences were downloaded from ENCODE. Archived files were ignored. We kept read-depth normalized signal files for DNase-seq and fold change over control files for ChIP-seq.

#### cCREs

cCREs for different biosamples were downloaded from SCREEN Registry V3 (https://screen.encodeproject.org/, by October 2021). cCRE count for each biosample ranges from 85248 to 138179 (Supplementary Table 1). DNase-only and Low-DNase elements were removed. All elements were padded to 350bp for the convenience of modeling, which is not mandatory.

#### Cell types

We selected human tissues, primary cells, cell lines, and *in vitro* differentiated cells 1) with RNA-seq, DNase, CTCF ChIP-seq, H3K4me3 ChIP-seq, and H3K27ac ChIP-seq data available on ENCODE and 2) with complete cCRE information on SCREEN Registry V3.

### CRE-gene interaction evaluation

#### Fulco et al

We downloaded the enhancer-gene interaction data from Supplementary Table 6a of the original study (13). We converted genomic coordinates of candidate enhancers from hg19 to hg38 using the liftover program of the UCSC Genome Browser (https://genome.ucsc.edu/cgi-bin/hgLiftOver). Non-autosomal genes were removed.

#### Gasperini et al

We downloaded the data from Table S2 of the original study (14). We converted genomic coordinates of candidate enhancers from hg19 to hg38 using the liftover program of the UCSC Genome Browser. To generate gene-mapped negative samples from the Gasperini dataset, we first selected target genes from the identified 664 enhancer-gene pairs, and then picked out candidate enhancers within the 1Mb region of each target gene respectively from all enhancers screened. Non-autosomal genes were removed.

#### Schraivogel et al

We downloaded the data from Supplementary Table 2 and 3 of the original study (15). We converted genomic coordinates of candidate enhancers from hg19 to hg38 using the liftover program of the UCSC Genome Browser. To generate gene-mapped negative samples from the Schraivogel dataset, we first selected target genes from the identified 41 enhancer-gene pairs, and then picked out candidate enhancers within the 1Mb region of each target gene respectively from all enhancers screened.

#### ChIA-PET

We obtained the ChIA-PET data of K562 from the ENCODE portal (ENCSR880DSH). To evaluate the model’s performance, for each gene, we used a total of 400 regulators upstream and downstream as the evaluation dataset. To calculate the cCRE-gene or cCRE-CRE interaction, for each pair of interacting sequences, we calculated whether the reads pair intersected the gene and CRE, respectively. The gene was considered to interact with the cCRE and regarded as a positive sample if crossed and as a negative sample otherwise.

#### ABC score

ABC score was adapted from Fulco et al(13). To be more specific, we collected the bigWig files of H3K27ac and DNase from ENCODE’s ENCFF977KGH and ENCFF414OGC, respectively, and converted them to bedGraph files with the UCSC tool `bigWigTobedGraph`. For each cCRE, we determined its signal by calculating the sum of the signals intersecting with it. Accordingly, we calculated the ABC score as the geometric mean of the H3K27ac and DNase signals multiplied by the reciprocal of the distance between the cCRE and the TSS (27).

### Classification of cCRE-gene interaction by distance groups

For each gene, the cCREs are divided into 4 groups (0-5kb, 5-50kb, 50-1000kb, 100-1000kb, 1000kb+) according to their distances to gene TSS. Groups with less than 10 gene-CREs pairs were filtered. auPRC and auROC for each group of each gene were calculated. For specificity and precision, we used mean values as the cutoff for the classification of positive and negative regulators.

### Enformer(27)

Pre-trained Enformer model was downloaded from https://github.com/deepmind/deepmind-research/tree/master/enformer. Genomic sequences flanking genes of interest were prepared following the original study’s instructions. Gradient x input of candidate enhancers was calculated following the original study’s instructions. To be pointed out, to simulate prediction tasks in new cell types, we used the cell-type-agnostic setting during the analysis. More specifically, the gradient was calculated and aggregated from all human tracks of the model.

### GraphReg(28)

Epi-GraphReg model was downloaded from https://github.com/karbalayghareh/GraphReg. Genomic sequences and DNase-seq, H3K27ac, H3K4me3 were prepared following the original study’s instructions. For a fair comparison, we incorporated histone modifications and transcription factor (TF) binding profiles used for CREaTor training as well (see Supplementary Table 4 for a full list). For training, we sourced chromosomes from cell lines including GM12878, B cells, HeLa-S3, MCF-7, fibroblast of dermis, CD14 positive monocyte, H1, HepG2, and keratinocyte (Supplementary Table 4c), deliberately excluding chromosomes Chr8, Chr9, and Chr16. The CAGE data for these cell lines were downloaded from ENCODE, as detailed in Supplementary Table 4c. We replaced the 3D genomic data with the reciprocal of the genomic distances between the cCRE and the TSS. For cCRE-gene interaction classification, we calculated saliency and integrated gradients for candidate enhancers following the original study’s instructions. The feature attribution type led to the best performance was used for comparison.

### TAD prediction

#### Hi-C data processing

We obtained the long-range chromatin interactions of K562Hi-C data from ENCODE (ENCSR545YBD). To estimate the interaction matrix with each cCRE as a bin, the Hi-C pairs that intersected with each cCRE pair were added together.

#### Calculation of insulation score

We calculated the sum of the interactions in each bin within 10kb as the Hi-C interaction matrix for 10kb resolution. A similar operation was applied to the attention matrix. We summed the min-max-normalized attention matrix within 10kb windows as the attention matrix at 10kb resolution. We obtained the location of the TAD boundary on K562 and GM12878 from the previous study (43). The interactions of the 3*3 matrix were summarized at one bin from the diagonal (58) to represent the insulation score for each TAD boundary. GM12878-specific TAD boundaries are genomics regions called in GM12878 boundary file exclusively.

#### Grouping of CTCF-bound elements

For all cCREs showing positive CTCF binding patterns, we determined whether they intersected with the TAD boundary from a previous study (43). We considered the intersecting cCREs as anchors of the TAD boundaries, and others as non-anchors. We extracted the attention scores between these CTCF-bound cCREs and then divided the weights into various groups. Scores for anchor cCRE pairs on the same TAD boundary were classified as “anchor-to-anchor”; scores between anchor cCRE and non-anchor cCRE within the same TAD were classified as “anchor-to-non-anchor”; scores for anchor cCREs on adjacent TADs were classified as “anchor-to-anchor in adjacent TADs”; and scores for anchor cCREs more than one TAD apart were classified as “anchor-to-anchor in remote TADs”.

### Mapping of CISH enhancers

First, we obtained the two regulatory loci Cp1 and Cp2 of CISH from the previous study(44) and converted their genomic coordinates from mm9 to mm10 using the `liftover` program of the UCSC Genome Browser. Then, for all cCREs of CISH genes in K562, we determined which cCREs intersected with Cp1 and Cp2, representing Cp1 and Cp2 respectively. Finally, we calculated the attention scores from Cp1 to Cp2 in K562 cell line to determine the effect of Cp1 on Cp2; and from Cp2 to Cp1 to determine the effect of Cp2 on Cp1. The control background (SDaCp1 and SDaCp2) consisted of interactions between cCREs with the same distance from Cp1 and Cp2 to CISH to all protein-coding genes except CISH.

### K562-specific genes

We obtained the expression data of each cell line’s gene from ENCODE (Supplementary Table 4a). Using the expression data of K562 as a control, we extracted the count matrix of each other cell line by the function `rsem-generate-data-matrix` of RSEM. These count matrices were then used to calculate differentially expressed genes using the function `rsem-run-ebseq`. After that, we screened the genes with PPDE (posterior probability that a gene/transcript is differentially expressed) greater than 95% as differentially expressed genes for K562 versus each cell line. Finally, the intersection of these differential genes was considered K562-specific genes.

### Representation clustering and visualization

First, we reduced the dimensionality of the 256-dimensional representations learned by our model with `scanpy.tl.pca` (default parameters). After a neighborhold graph is calculated (`scanpy.pp.neighbors`, `n_neighbors=20`, `n_pcs=50`), we clustered reduced representations with Leiden graph-clustering method (`scanpy.tl.leiden`, `resolution=0.5`). The neighborhood graph and clusters were then visualized using Uniform Manifold Approximation and Projection (UMAP).

## Data Availability

RNA expression, DNase-seq, ChIP-seq, Hi-C, CAGE and ChIA-PET files were downloaded from https://www.encodeproject.org/ (Supplementary Table 4). cCREs for different biosamples were downloaded from SCREEN Registry V3 (https://screen.encodeproject.org/). Both K562 TAD boundary and GM12878 TAD boundary file were downloaded from https://drive.google.com/drive/folders/15Rc6PhrrBjThwE-5dSyNX-ILELaUu6uG. CRISPR perturbation experiments of enhancer-gene interactions were downloaded from reference 13-15 respectively.

## Code Availability

The code for data processing, model training and evaluation are available at https://github.com/DLS5-Omics/CREaTor.

## Acknowledgements

We thank Dr. Chuan Cao, Dr. Haiguang Liu, Dr. Bofeng Liu, Dr. Han Yuan, and Dr. Yanxiao Zhang for constructive suggestions and feedback; Dr. Tie-Yan Liu for project supervision; Shizhuo Zhang and Yifan Deng for discussions on modeling.

## Authors’ contributions

Conceptualization, P.D., Y.L., H.X. and Y.Q.; methodology and modeling, F.J., Z.C., P.D., H.X. and Y.Q.; data curation, P.D., Y.L., F.J. and Z.C.; result interpretation, P.D., Y.L., F.J., and Y.Q.; writing-original draft, P.D., Y.L., Z.C., Y.Q, and F.J.; writing-review, P.D., Y.L., Y.Q, H.X., Z.C., L.H., L.W., F.J., B.S. and J.Z.; supervision, B.S.. All authors read and approved the final manuscript.

---

**Extended Data Figure 1.**
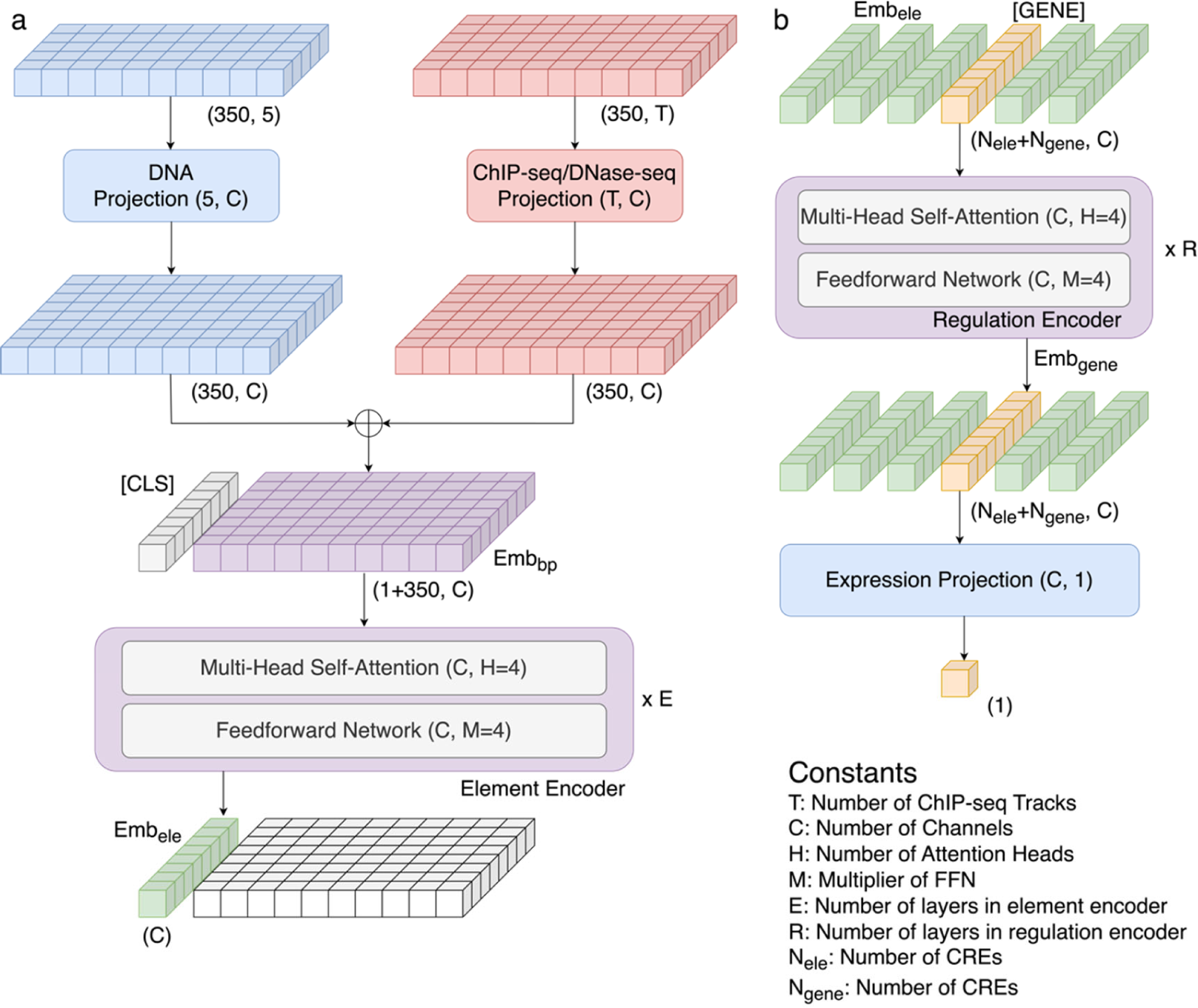
Architecture of CREaTor. CREaTor is composed of two modules. **a)** Element module encodes the representation of cCREs. We first map DNA and ChIP-seq/DNase-seq to latent space through a linear projection respectively, and then combine them through element-wise addition to obtain *Emb_bp_*, the feature embedding of each bp. We feed the *Emb_bp_*s into the element encoder together with a [CLS] token. The [CLS] token adaptively aggregates information from the *Emb_bp_*s in the element encoder. We use the output vector of [CLS] token as the representation of the element, namely *Emb_ele_*. **b)** Regulation module models the interaction between cCREs and genes. We concatenate the *Emb_ele_* of cCREs (denoted in blue and yellow) and the [GENE] tokens (denoted in red) as the input of the regulation encoder. The [GENE] tokens interact with and are regulated by the cCREs in the regulation encoder. We apply a linear projection with SoftPlus activation on the output vector of [GENE] tokens to predict their expressions. The size of each component of the architecture is shown as a tuple inside the block. The shape of the tensor at each step is denoted as a tuple in the bottom right of the blocks.

**Extended Data Figure 2.**
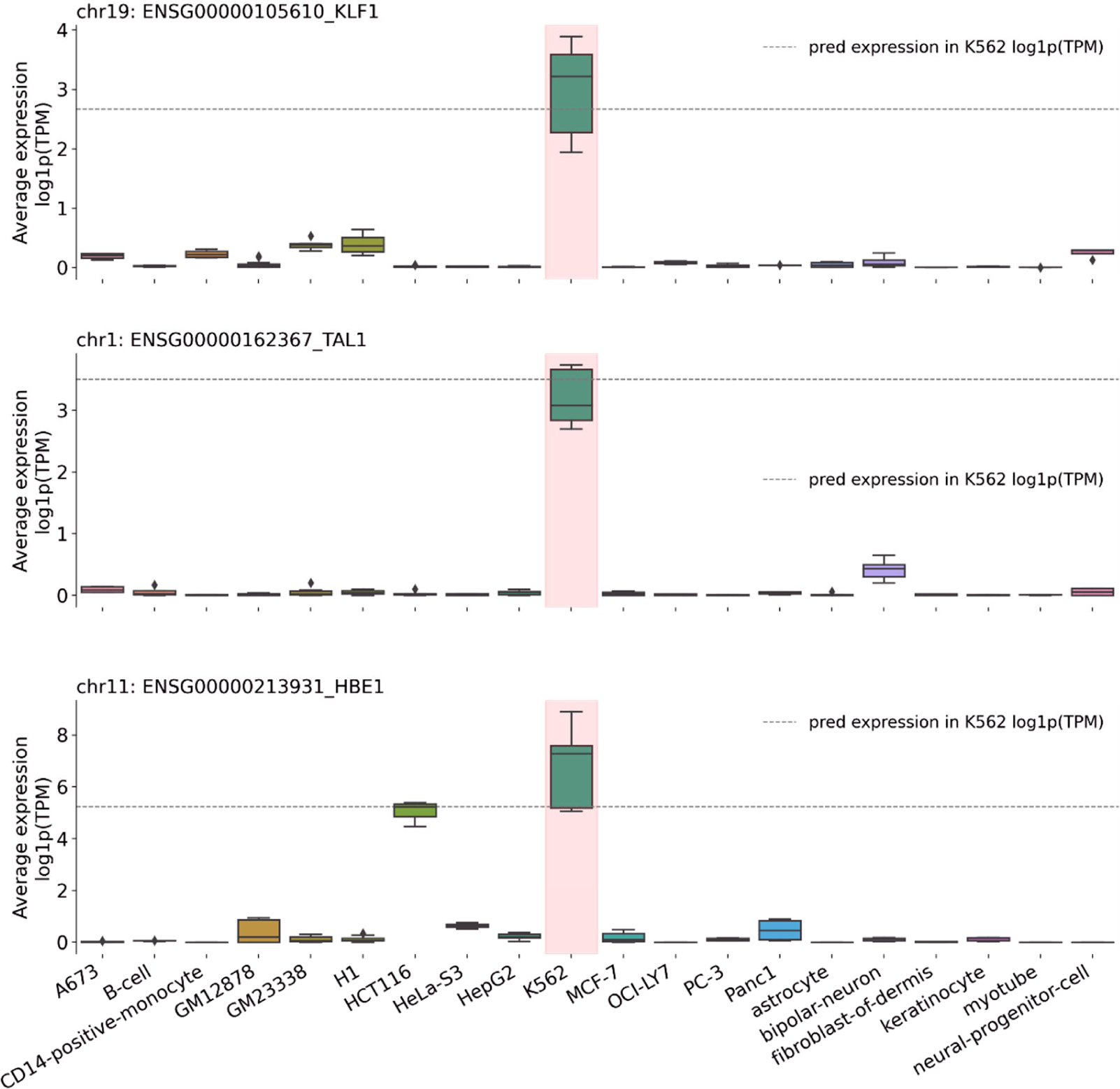
Prediction of K562 differentially expressed genes. Representative examples of observed and predicted expression of genes KLF1, TAL1 and HBE1 in 20 different types of cells. The dashed line indicates the predicted values.

**Extended Data Figure 3.**
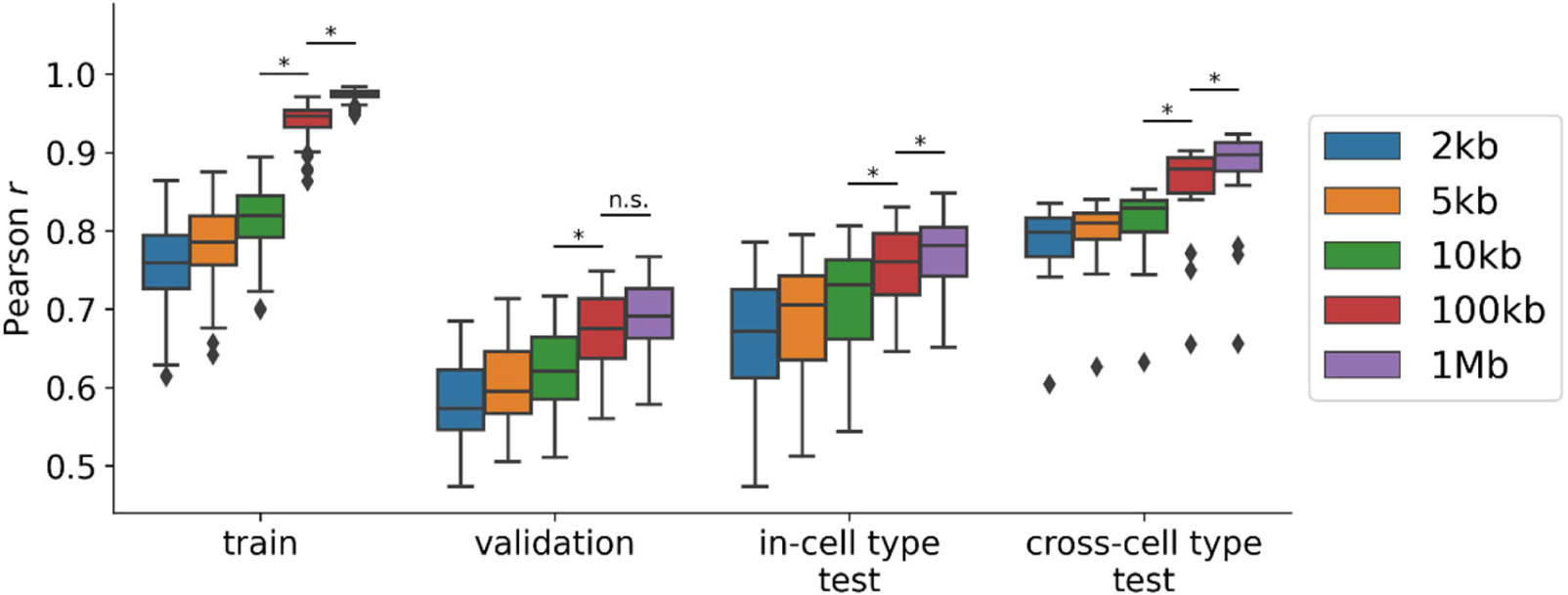
Performance of CREaTor with cCREs up to 2kb, 5kb, 10kb, 100kb, or 1Mb away from the TSS of target genes. The training set includes chr1-7, 10-15, and 17-22 in 19 cell types other than K562. The validation set includes chr16 in cell types other than K562. The in-cell type test set includes chr8 and chr9 in cell types other than K562. Cross-cell type test set represents all chromosomes in the K562 cell line. P values were computed with the two-sided Mann–Whitney U test.

**Extended Data Figure 4.**
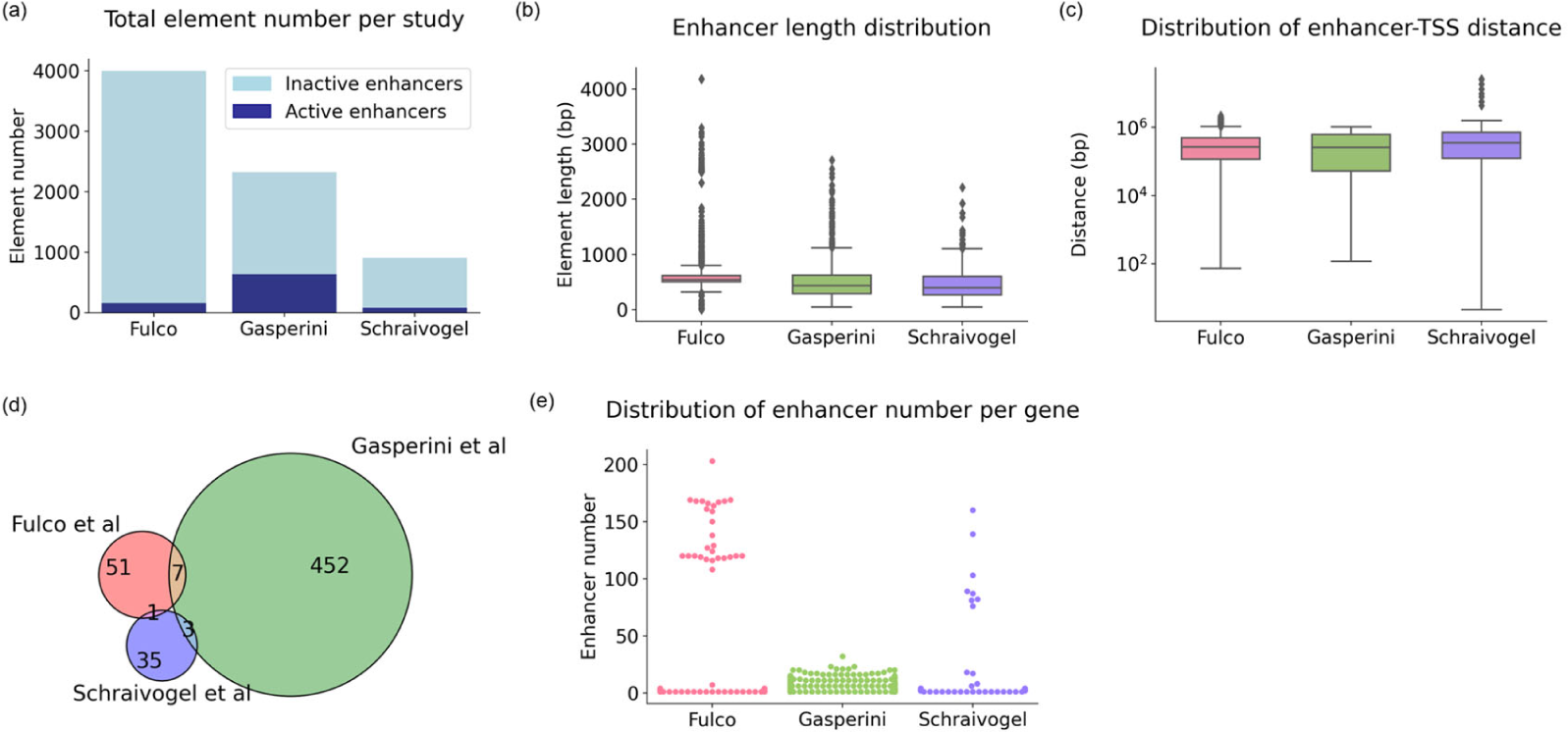
Statistics of enhancer-gene interaction data from 3 CRISPRi-based studies. The statistics were performed on data after genomic coordinates liftover and non-autosomosal data filtering. (a) The number of active and inactive enhancers tested by each study. (b) Enhancer length distribution in each study. (c) Enhancer-gene TSS distance distribution in each study. (d) Overlapped active enhancers in 3 studies. (e) The number of enhancers tested for each gene in each study.

**Extended Data Figure 5.**
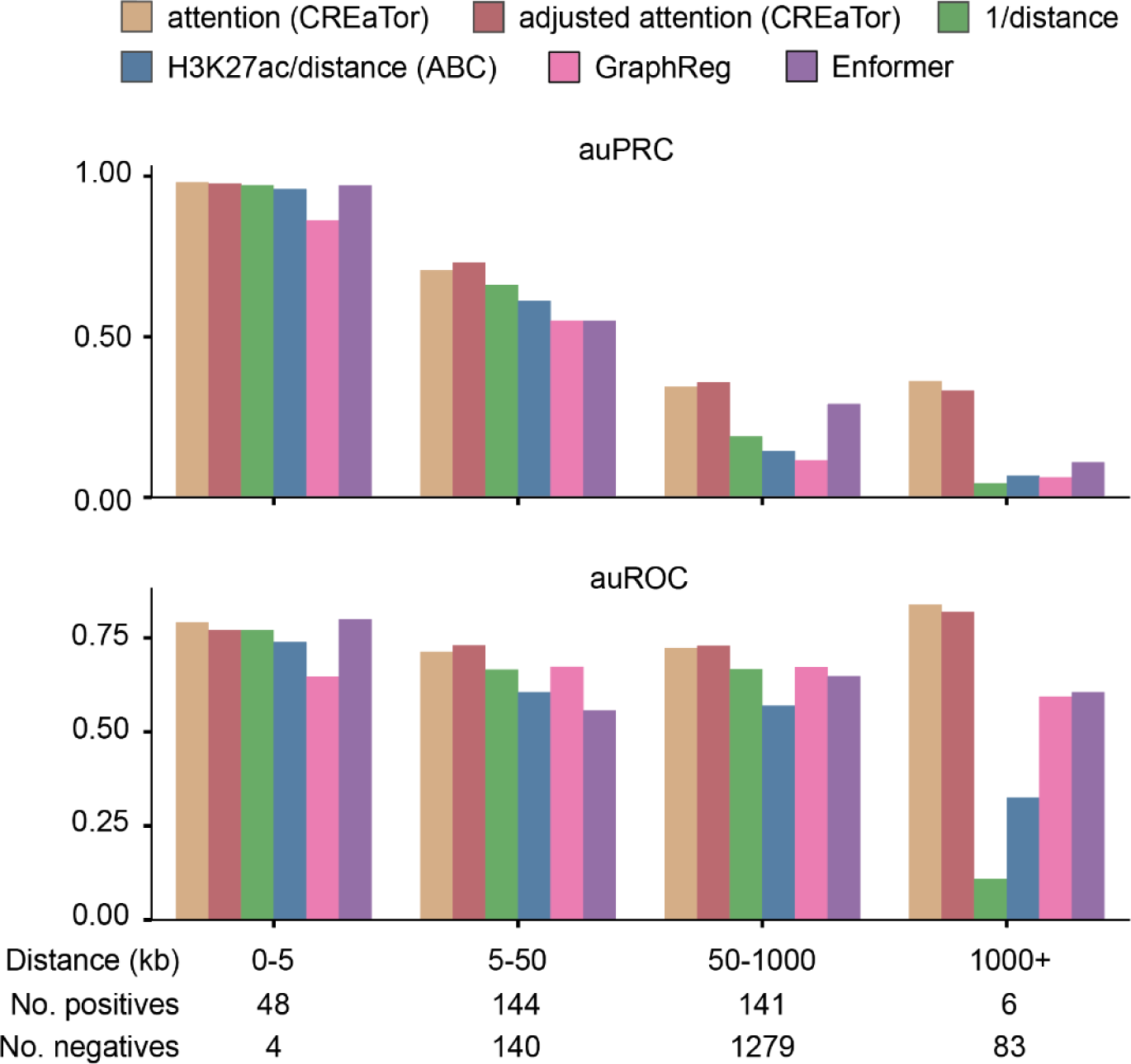
auPRC and auROC of CREaTor and its counterparts on the classification of cCRE-gene pairs collected from 3 independent CRISPR perturbation experiments. cCRE-gene pairs are stratified by their relative genomics distances. The number of positive/negative labels in each group is annotated at the bottom. Labels (positive/negative) of cCRE-gene pairs were collected from CRISPR perturbation experiments.

**Extended Data Figure 6.**
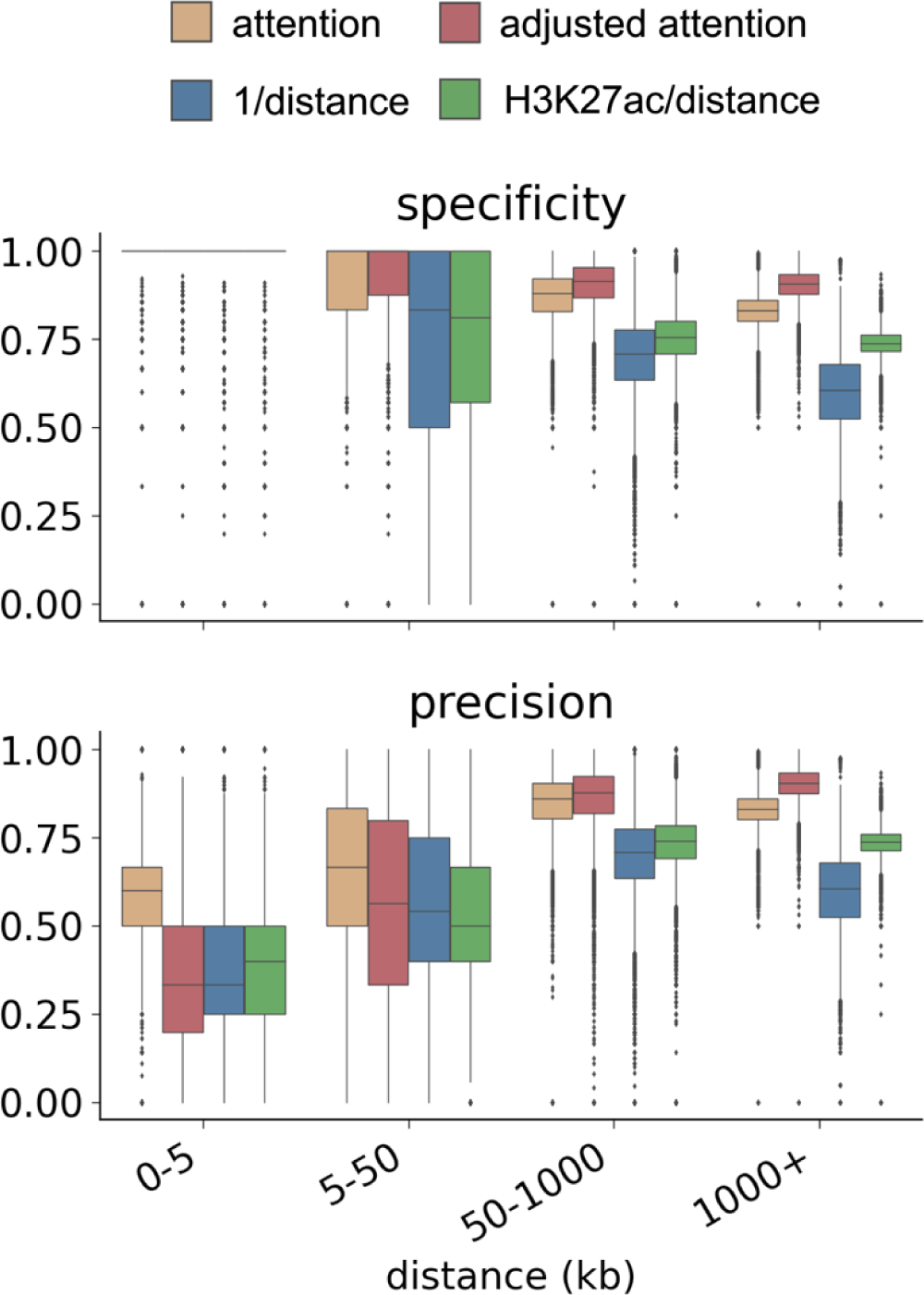
Specificity and precision scores of CREaTor and its counterparts on cCRE-gene pair classification. Distance denotes the relative genomic distance between cCREs and genes. The performance is evaluated for each gene and each distance group separately. The H3K27ac value of a cCRE is calculated as the sum of the H3K27ac peak values of the element. Positive/negative cutoff is set as mean values of attention scores in each distance group. Labels (positive/negative) of cCRE-gene pairs were collected from a Pol-II mediated ChIA-PET experiment of K562.

**Extended Data Figure 7.**
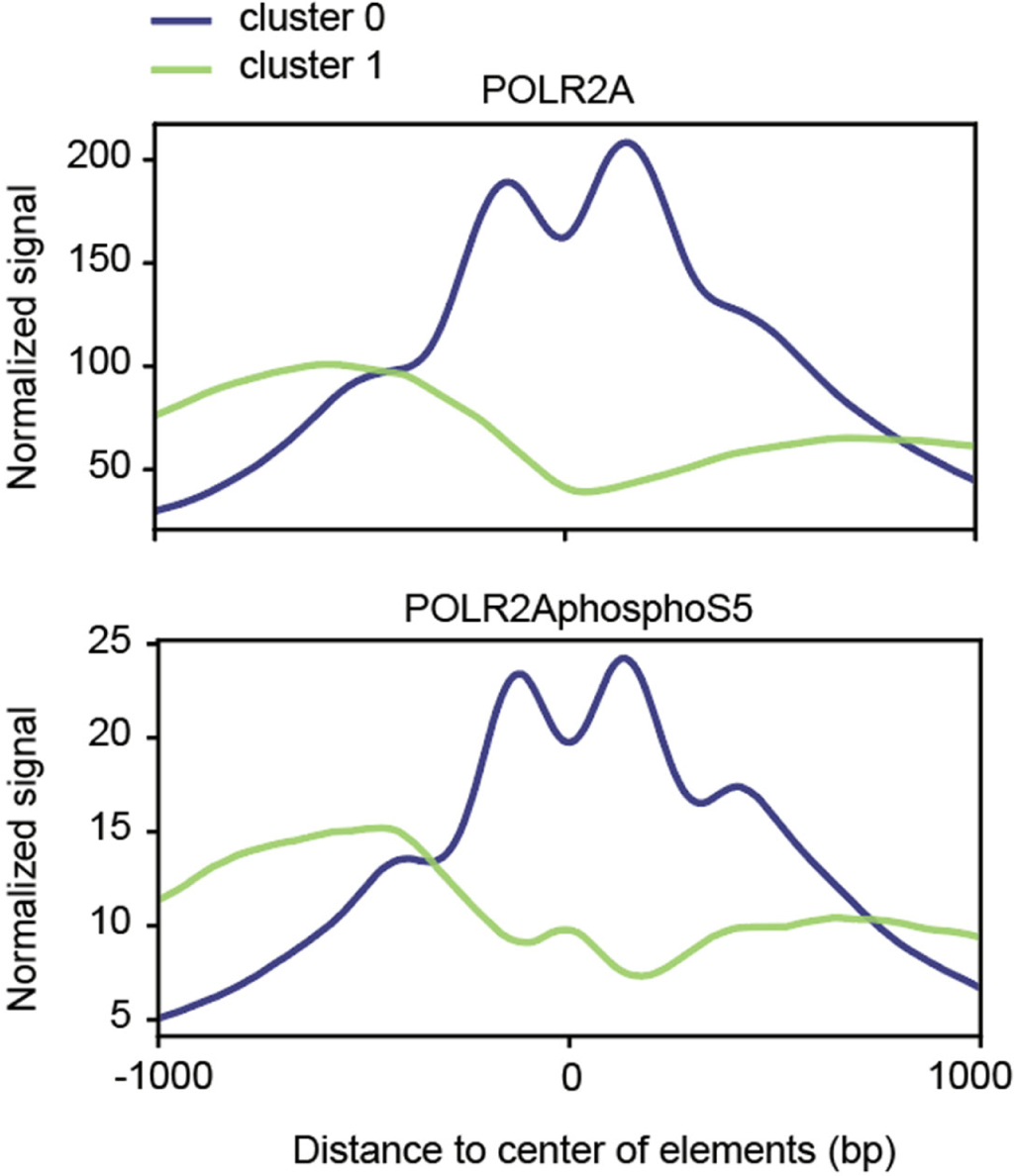
Average signals of RNA Pol II on cCREs in cluster 0 and cluster 1 respectively. Upper: unphosphorylated form of Pol II. Bottom: Pol II CTD phospho Ser5.

**Extended Data Figure 8.**
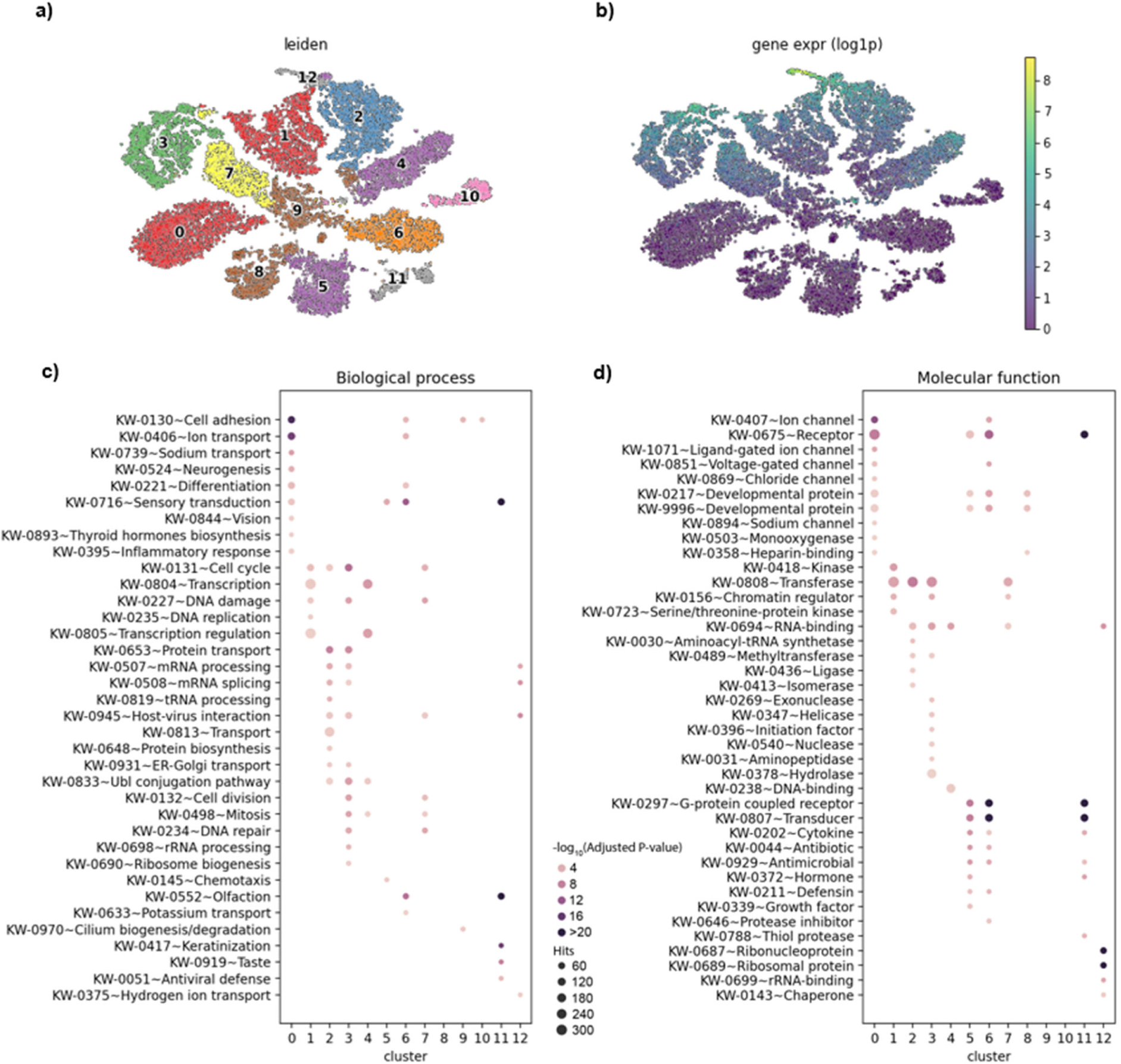
Gene representations learned by CREaTor can be clustered into groups with different functions. **a)** Uniform Manifold Approximation and Projection (UMAP) of gene embedding in K562, colored and numbered as clusters grouped by the Leiden algorithm. **b)** Same as (a), but colored by gene expression levels. **c)** Functional annotation clustering with the DAVID Gene Functional Classification Tool (DAVID, http://david.abcc.ncifcrf.gov) using UniProtKB biological process keywords. Significantly enriched (adjusted p-value<0.05) groups for genes in each cluster in (a) are shown. **d)** Functional annotation clustering with DAVID using UniProtKB molecular function keywords. Significantly enriched (adjusted p-value<0.05) groups for genes in each cluster in (a) are shown.

