## Supplementary Figures for "CREaTor: Zero-shot *cis*-regulatory pattern modeling with attention mechanisms"

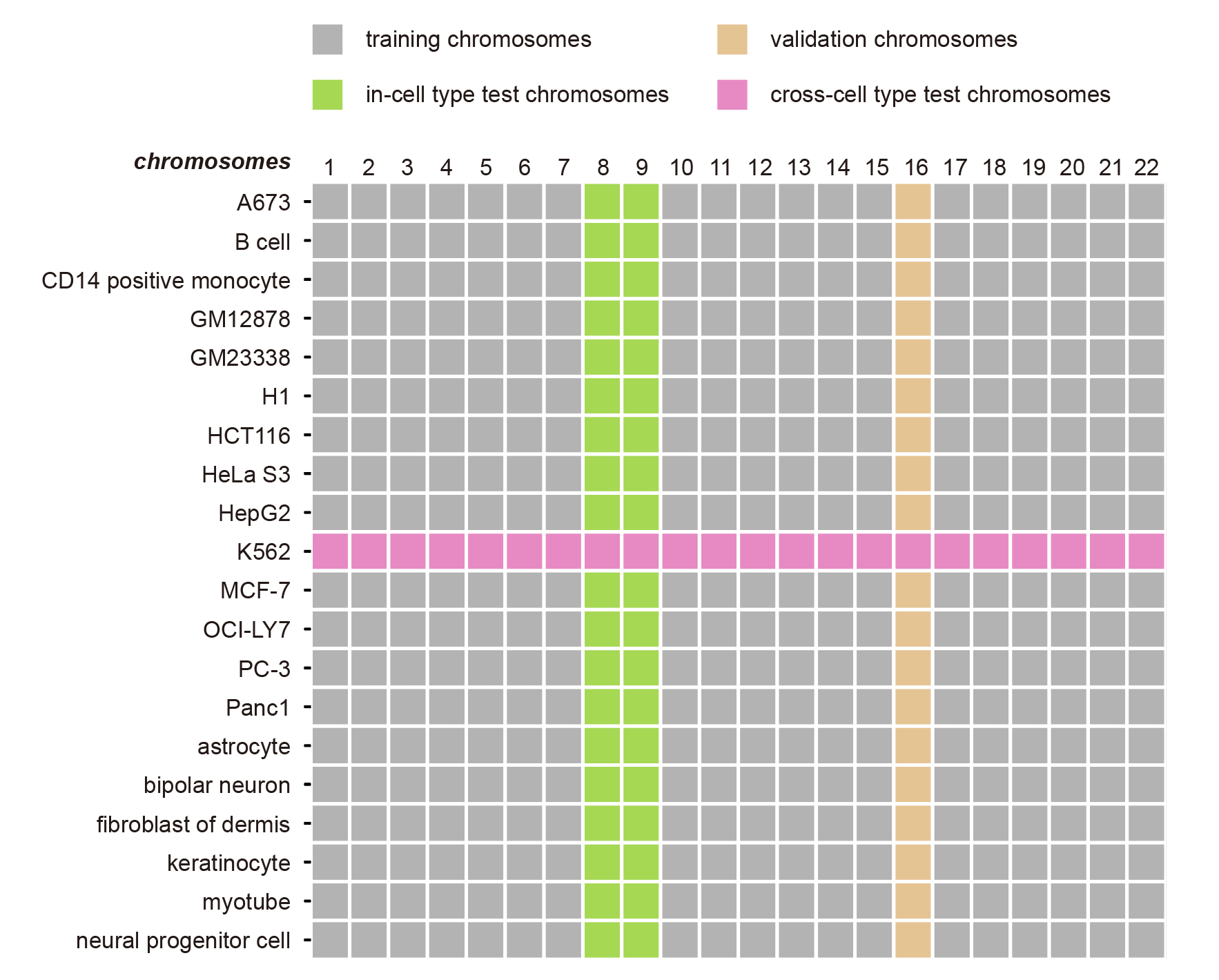

**Supplementary Figure 1.** **Data split strategy for CREaTor Training and evaluation.** Training chromosomes (grey): chr1-7, chr10-15, and chr17-23 of 19 human tissues and cell lines: A673, B cell, CD14 positive monocyte, GM12878, GM23338, H1, HCT116, HeLa S3, HepG2, MCF-7, OCI-LY7, PC-3, Panc1, astrocyte, bipolar neuron, fibroblast of dermis, keratinocyte, myotube, and neural progenitor cell. Validation chromosomes (khaki): chr16 of the same 19 human tissues and cell lines. Additionally, we designed 2 sets of test datasets: In-cell type test chromosomes (green) denote chr8 and chr9 in the 19 human tissue and cell lines (excluding K562). Cross-cell type test chromosomes (magenta) denote all autosomes of K562 cell line, as K562 is not used for model training.

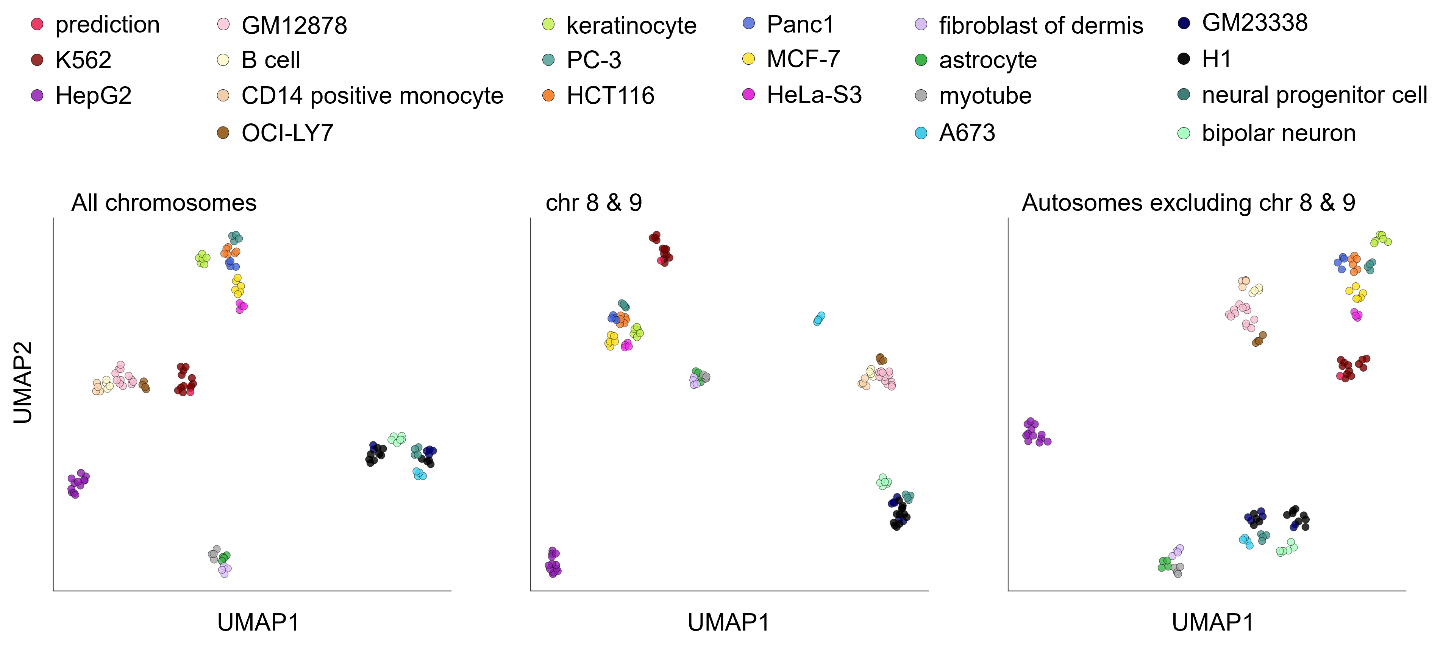

**Supplementary Figure 2.** **UMAP visualization of 123 expression profiles.** Prediction: Predicted K562 gene expression profile. Others: Transcript quantifications of corresponding cell types with RNA-seq.

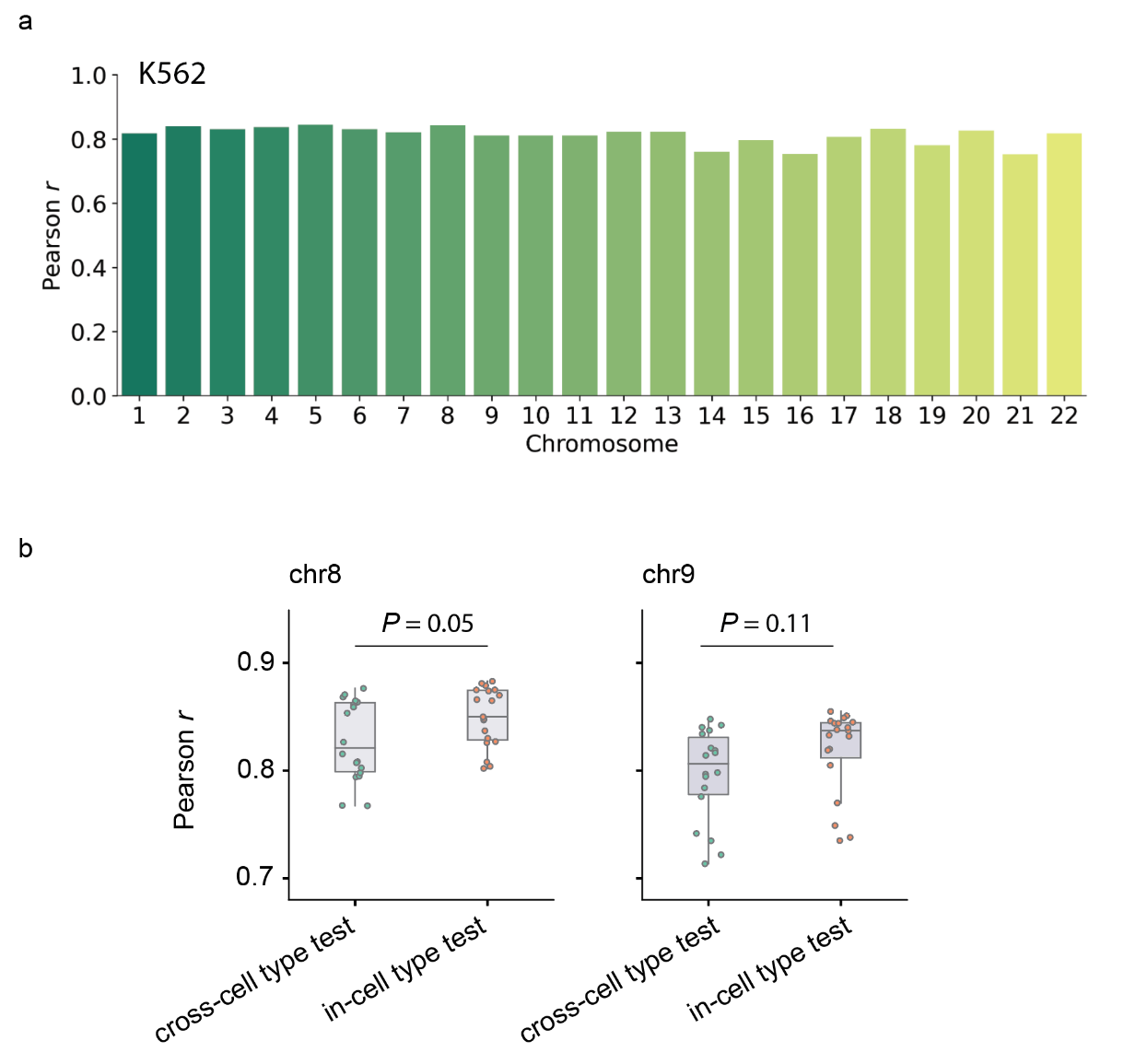

**Supplementary Figure 3.** **Leave-one-chromosome-out and leave-one-cell type out assays.** **a)** Pearson *r* between predicted and observed expression of genes on chr1-22 in K562. In this leave-one-chromosome-out assay, each chromosome was excluded from training once, and the corresponding chromosome in K562 was used for test (Supplementary Table 2a). **b)** Pearson *r* between predicted and observed expression of genes on chr8 and chr9 in different cell types under different settings. Cross-cell type test: as in the leave-one-cell type-out setting, all chromosomes of each cell type were held out for test once (Supplementary Table 2b). In-cell type test: The cell type was used for modeling training, but chr8-9 were held out for test (Extended Data Table 1). Each dot represents one cell type. P values were computed with the two-sided Mann–Whitney U test.
